## Supplementary Figures for "Combinatorial multimer staining and spectral flow cytometry facilitate quantification and characterization of polysaccharide-specific B cell immunity"

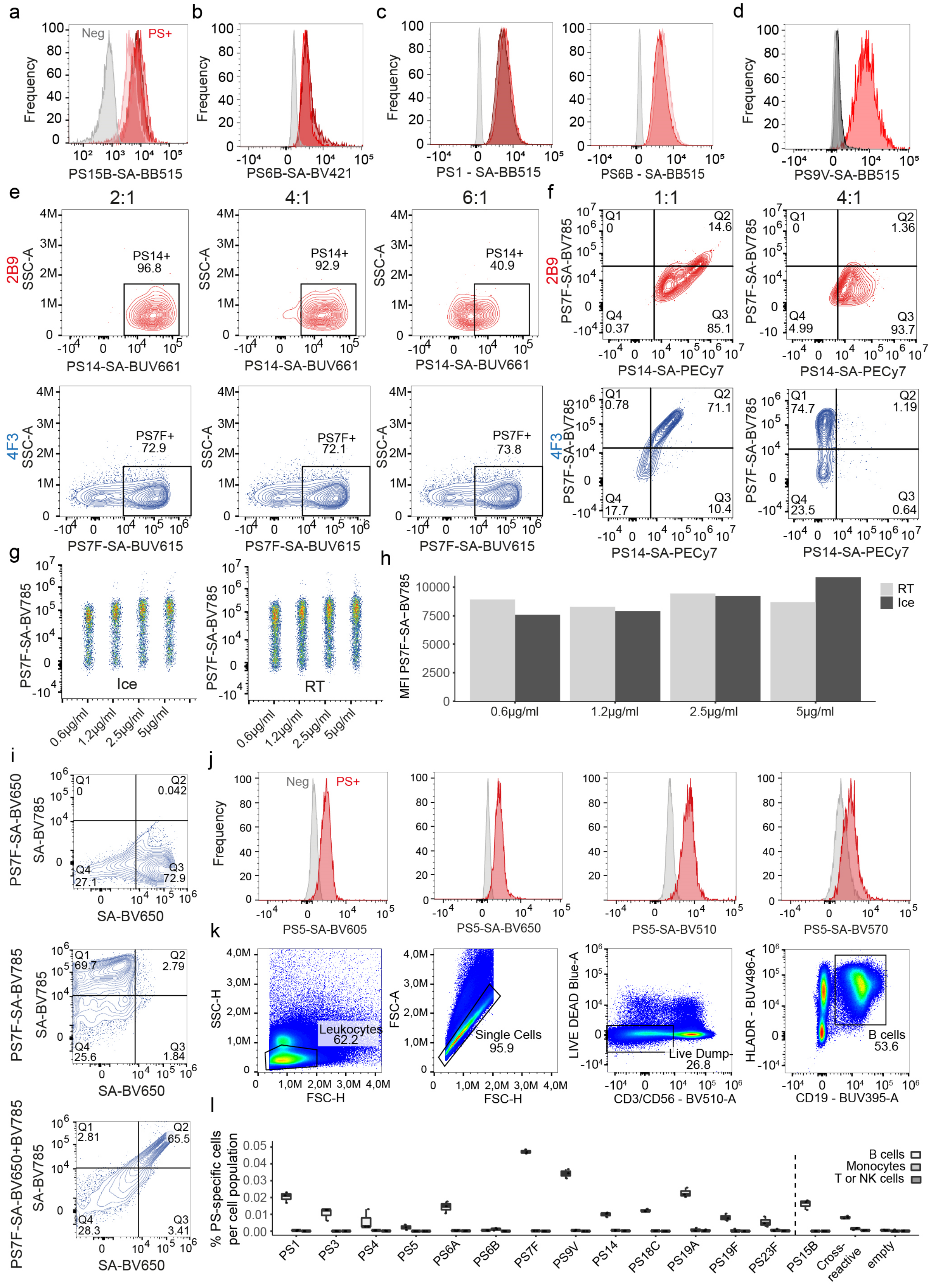

**Supplementary Fig. 1. Optimization and validation of assay conditions. a**, Histogram of four batches of PS15B-SA-BB515 multimers (PS+; shades of red) prepared on different 4 days and recorded in 2 different readings, showing similar signal intensity, normalised to mode. Control unstained beads are depicted in grey (Neg). **b**, Signal of PS6B-SA-BV421 on specific antisera-covered compensation beads, normalised to mode, using multimers made from the same batch of biotinylated PS6B, either kept in at 4$^{o}$C (dark red) or -80$^{o}$C (light red) and multimerized with SA-BV421 three months later. **c**, Histograms of PS-SA-BB515 multimers, using freshly biotinylated (light red) PS1 (left panel) and PS6B (right panel) or biotinylated PS stored at -80$^{o}$C for 8-12 months (dark red), normalised to mode. Histograms include negative beads (grey). **d**, Specificity of antisera shown by signal of PS9V-SA-BB515 on compensation beads coupled to autologous antiserum pool R (red), heterologous antiserum factor 6C (black), or without antiserum (grey), normalised to mode. **e**, Spectral flow cytometry density plots showing 2B9 (red; PS14-specific) and 4F3 (blue; PS7F-specific) clones binding PS7F-SA-BV785 or PS14-SA-PECy7 multimers prepared with a PS:SA ratio of 2:1, 4:1 or 6:1, respectively. **f**, Spectral flow cytometry results of clone 2B9 (red; PS14-specific) and 4F3 (blue; PS7F-specific) stained simultaneously with PS7F-SA-BV785 and PS14-SA-PECy7 multimers, prepared with a PS:SA ratio of 1:1 or 4:1. The top right quadrant indicates double positive cells, which points to non-specific binding. **g**, Density plots of 4F3 (PS7F-specific) clones stained with various concentrations of PS7F-SA-BV785 multimers on ice for 30 minutes or at room temperature (RT) for 15 minutes. **h**, Bar plots showing the multimer MFI signal from PBMCs of a donor with no known PCV13 vaccination, stained with various concentrations of PS7F-SA-BV785 multimers on ice (30 minutes) or at room temperature (RT; 15 minutes). **i**, Spectral flow cytometry density plots showing 4F3 (PS7F-specific) clones stained with PS7F-SA-BV650 (top panel), PS7F-SA-BV785 (middle panel) or both simultaneously (bottom panel). **j**, Signal of discarded fluorochromes. Biotinylated PS5 in multimer with SA conjugated to different fluorochromes were bound to compensation beads using PS-specific antiserum (PS+; red), normalised to mode. Negative beads are shown for control (Neg; grey). **k**, Gating strategy of PBMCs stained with the panel indicated in (Supplementary table 1). **l**, Frequencies of PS-specific cells in B cells ($SSC^{low}CD3^{-}CD56^{-}CD19^{+}HLADR^{+}$, light grey), monocytes ($SSC^{high}CD3^{-}CD56^{-}CD19^{-}$, darker grey) or T or NK cells ($SSC^{low}CD3^{+}CD56^{+}CD19^{-}$, darkest shade of grey) in PBMCs from a donor 3 weeks post-PCV13 vaccination.

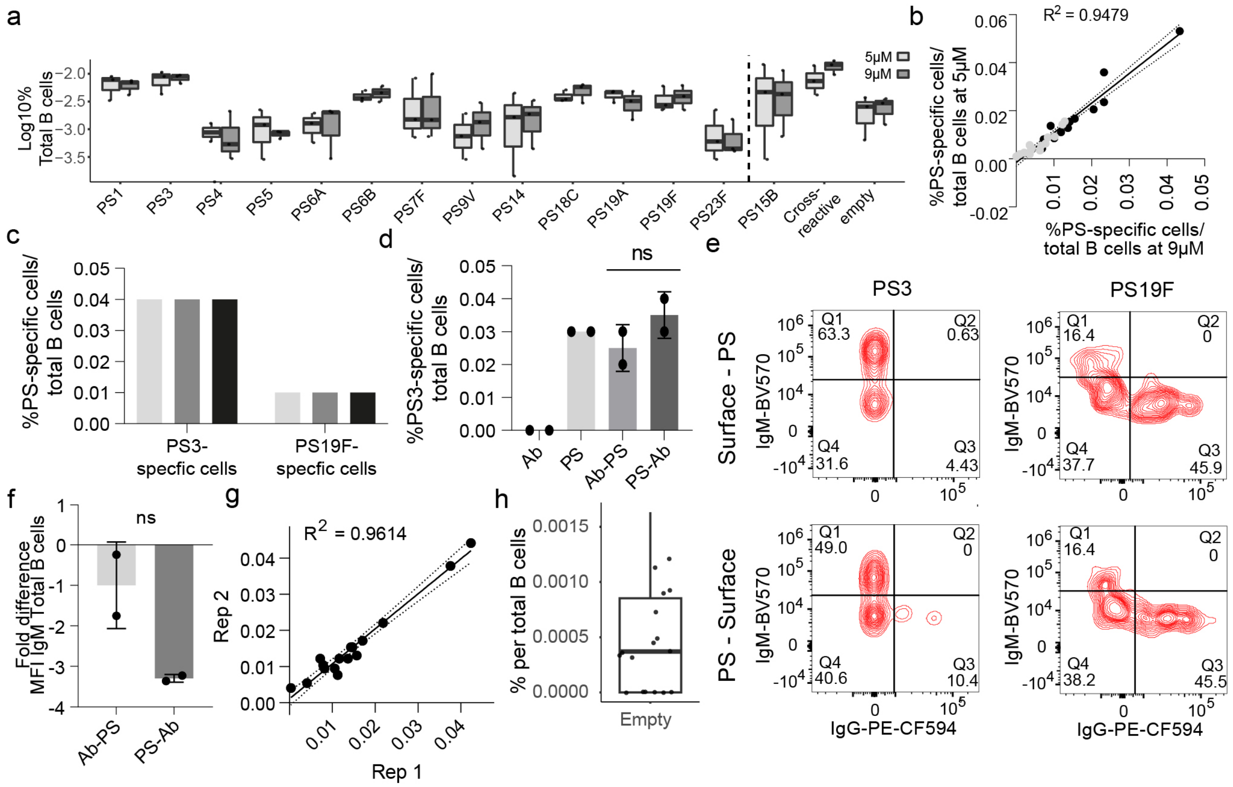


**Supplementary Fig. 2. Validation of PBMC probing with PS-SA multimers. a**, Frequency of PS-specific cells in PBMCs from 3 donors with assumed no PCV13 vaccination, stained with 5µM (light grey) or 9µM (dark grey) PS-SA multimers for 14 serotypes. **b**, Correlation plot showing similarity between the staining of PBMCs with different concentrations in (a). Statistical analysis by Pearson correlation. **c**, Frequencies of PS3- and PS19F-specific B cells in PBMCs from a donor 3 weeks post-PCV13 vaccination stained with three different SA-fluorochrome combinations on PS3 and PS19F, namely BB515/BV711 (light grey), BUV615/BV421 (dark grey) and BUV5661/BV785 (black). **d**, Frequency of PS3-specific B cells in PBMCs from a donor 3 weeks post-PCV13 vaccination stained with only surface marker antibodies (Ab) or PS-SA multimers (PS), stained first for surface markers and subsequently with PS-SA multimers (Ab-PS) or stained first with PS-SA multimers and for surface markers afterwards (PS-Ab). Average and standard deviation shown of two technical duplicates. Statistical analysis by Mann-Whitney test (P=0.6667). **e**, Contour plots showing IgM and IgG MFI and positive population frequencies on PS3- and PS19F-specific cells from (d). **f**, Fold-difference of IgM MFI between total B cells from surface staining only group compared to Ab-PS and PS-Ab from (d). Statistical analysis by Mann-Whitney test (P=0.3333). **g**, Correlation between two experimental duplicates of stained PBMCs from one donor 3 weeks post-PCV13. The black lines indicates linear regression ($R^2$=0.9614) and the 95% confidence interval. Statistical analysis by Pearson correlation. **h**, Frequency of cells in the "empty" category ($BUV615^+BV711^+$) from all donors from (Fig. 3) to indicate the background and lower limit of detection. Average, the standard deviation and individual results of three independent experiments are shown.

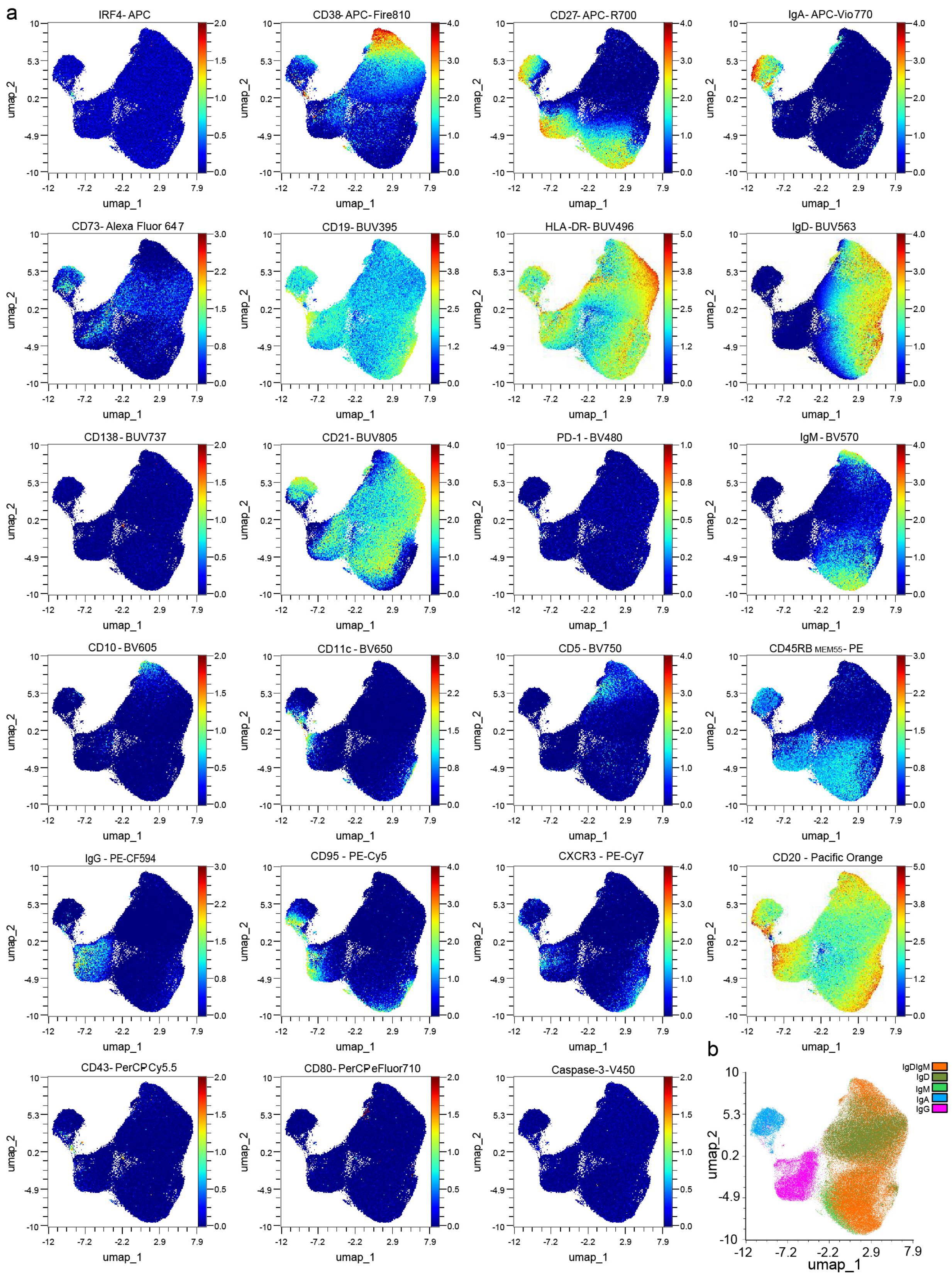
a
IRF4- APC
CD38- APC- Fire810
CD27- APC- R700
IgA- APC-Vio770
CD73- Alexa Fluor 647
CD19- BUV395
HLA-DR- BUV496
IgD- BUV563
CD138 - BUV737
CD21- BUV805
PD-1 - BV480
IgM - BV570
CD10 - BV605
CD11c - BV650
CD5 - BV750
CD45RB MEM55 - PE
IgG - PE-CF594
CD95 - PE-Cy5
CXCR3 - PE-Cy7
CD20 - Pacific Orange
CD43- PerCPCy5.5
CD80- PerCPeFluor710
Caspase-3-V450
umap_1
umap_2
b
IgDIgM
IgD
IgM
IgA
IgG

**Supplementary Fig. 3. Marker expression distribution across the UMAP. a**, Density plots from all clustered B cells from (Fig. 4). **b**, Clusters of $IgD^{+}IgM^{+}$ (orange), $IgD^{+}$ (dark green), $IgM^{+}$ (light green), $IgA^{+}$ (blue) and $IgG^{+}$ (magenta) gated B cells on a UMAP, from (Fig. 4a-b).

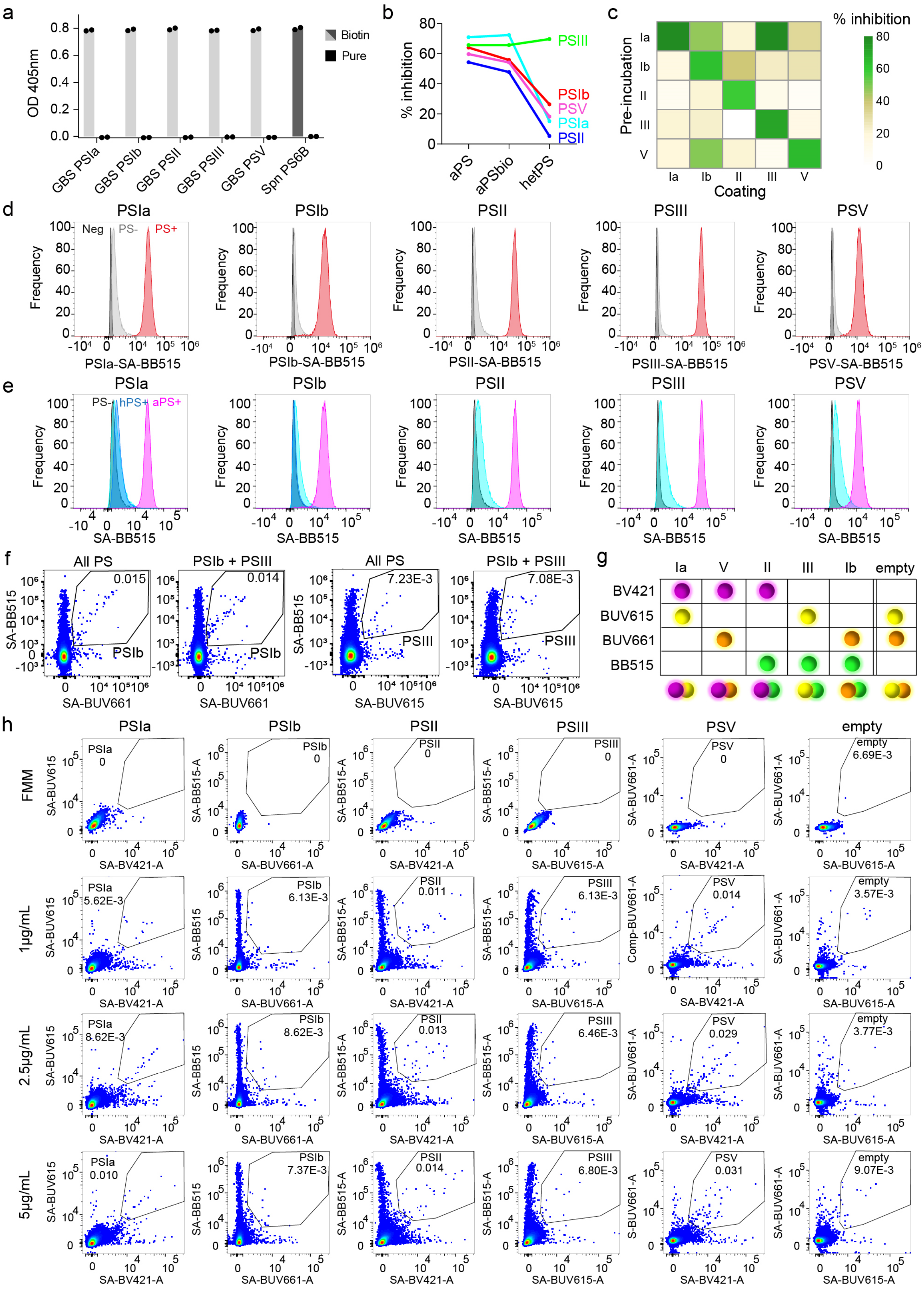
a
b
c
d
e
f
g
h
OD 405nm
Biotin
Pure
GBS PSIa
GBS PSIb
GBS PSII
GBS PSIII
GBS PSV
Spn PS6B
% inhibition
aPS
aPSbio
hetPS
PSIII
PSIb
PSV
PSIa
PSII
Pre-incubation
Coating
Ia
Ib
II
III
V
Frequency
Neg
PS-
PS+
PSIa-SA-BB515
PSIb-SA-BB515
PSII-SA-BB515
PSIII-SA-BB515
PSV-SA-BB515
hPS+
aPS+
SA-BB515
All PS
PSIb + PSIII
0.015
0.014
7.23E-3
7.08E-3
SA-BUV661
SA-BUV615
empty
BV421
BUV615
BUV661
BB515
FMM
1μg/mL
2.5μg/mL
5μg/mL
SA-BV421-A
SA-BUV661-A
SA-BUV615-A
SA-BB515-A
SA-BUV661-A
Comp-BUV661-A
6.69E-3
5.62E-3
6.13E-3
0.011
0.014
3.57E-3
8.62E-3
0.013
6.46E-3
0.029
3.77E-3
0.010
7.37E-3
6.80E-3
0.031
9.07E-3

**Supplementary Fig. 4. Optimization of GBS PS biotinylation and PS-multimer formation. a**, Bar plots with biotin ELISA results showing the efficacy of biotin incorporation in CDAP-activated GBS PS (light grey) and pneumococcal PS6B as positive control (dark grey). Untreated PS are shown as negative control (black). Blank-corrected OD values are shown. For each PS, levels were measured in duplicate, indicated by symbols with average depicted by bars. **b**, Competition ELISA showing the percentage of antibody blocking by biotinylated and non-modified PS. For each PS, inhibition of detection by preincubation of pooled donor samples (confirmed presence of serotype-specific antibodies), with autologous unmodified PS (aPS), autologous biotinylated PS (aPSbio), heterologous unmodified PS (hetPS) is shown. Inhibition indicates the reduction in OD signal in pre-absorbed versus non-pre-absorbed serum. Average of duplicates is shown. **c**, Competition ELISA as (b), showing all autologous and heterologous combinations of non-biotinylated PS, to show cross-reactivity of serotype-specific antibodies. Average of duplicates is shown. **d**, Histograms showing fluorescent signal of PS-SA multimers on compensation beads alone (Neg; black), with PS-SA-multimers (PS-; grey) or beads coupled with PS-specific autologous antiserum and stained with PS-SA-multimers (PS+; red), normalised to mode. **e**, Histograms showing signal of compensation beads with PS-SA-multimers alone without antiserum to indicate background (PS-; black) or compensation beads coupled to autologous serotype-specific antiserum and stained with autologous (aPS+; magenta) or heterologous (hPS+; light blue are combined heterologous PS, dark blue are heterologous PSIa/PSIb) PS-SA-multimers, normalised to mode. **f**, Frequency of PSIb and PSIII-specific cells in pooled PBMCs from 6 South African donors, after staining for all 5 serotypes (PSIa, PSIb, PSII, PSIII and PSV) or solely 2 (PSIb and PSIII). **g**, Barcode method of combinatorial staining patterns for GBS PS-SA multimers. The four fluorochromes are indicated in rows and the various serotypes in columns, including an empty colour combination for background control. **h**, Staining of pooled PBMCs of South African donors, with viability dye, CD3, CD19 and 3 concentrations of PS-SA-multimers: 1µg/mL, 2.5µg/mL and 5µg/mL. The fluorescence minus tetramer (FMM) control sample was made by pooling all samples and staining with everything but tetramers.

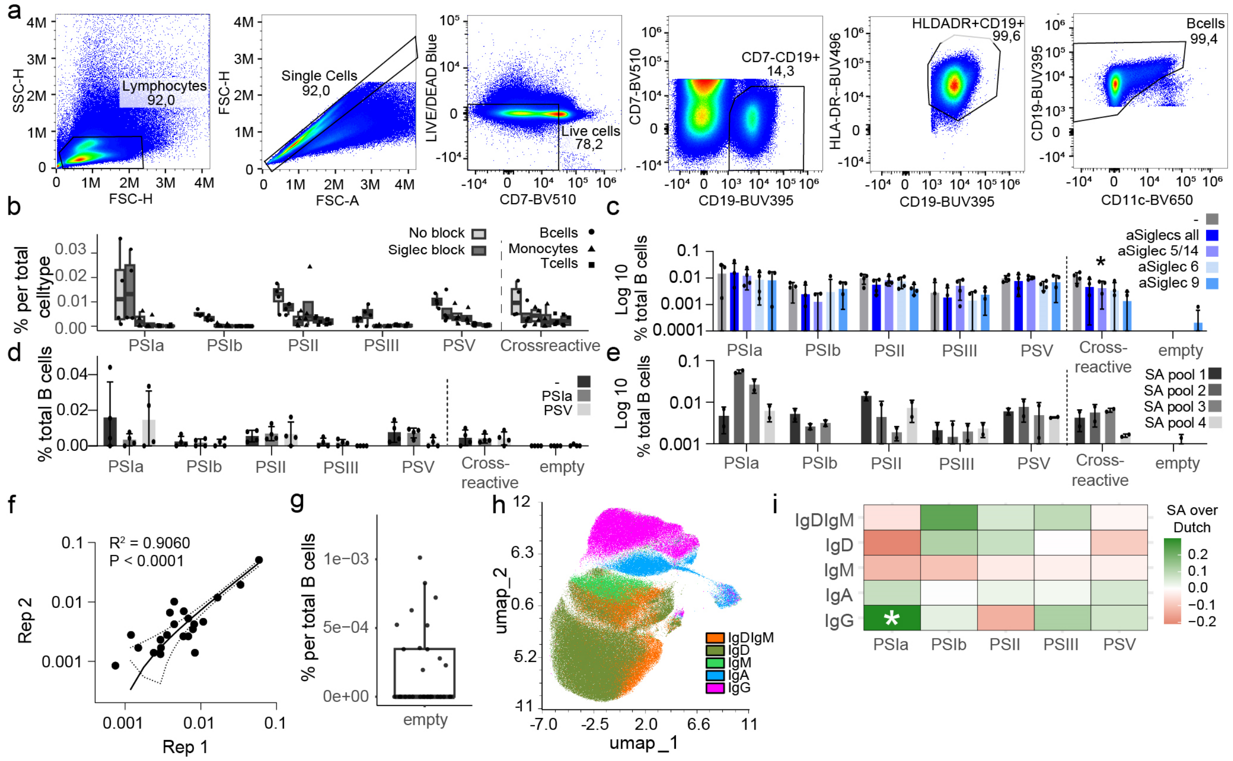


**Supplementary Fig. 5. Verification on GBS PS-multimer staining.**

**a**, Gating strategy for Dutch and South African PBMCs stained with the panel in (Supplementary table 4). **b**, Frequencies of PS-specific cells in B cells (SSC$^{low}$CD3$^{-}$CD19$^{+}$HLADR$^{+}$, circles), monocytes (SSC$^{high}$CD3$^{-}$CD19$^{-}$, triangles) and T cells (SSC$^{low}$CD3$^{+}$CD19$^{-}$, squares), in pooled PBMCs from South African donors. Siglec receptors were either blocked (dark grey) with 10µg/mL each of anti-Siglec-5/14 (RnD systems, MAB10721-SP), anti-Siglec-6 (RnD systems, MAB2859-SP), anti-Siglec-9 (RnD systems, MAB1139-100), or non-blocked (light grey). Average and standard deviation are shown. **c**, Signal of pooled PBMCs of South African donors pre-blocked (shades of blue) with 10µg/mL anti-Siglec-5/14 (RnD systems, MAB10721-SP), anti-Siglec-6 (RnD systems, MAB2859-SP), anti-Siglec-9 (RnD systems, MAB1139-100), 30µg/mL anti-Siglec all combined (10µg/mL each) or nothing (grey), and stained with viability dye, CD3, CD19 and PS-SA multimers. **d**, Frequency of PS-specific B cells in pooled PBMCs from South African donors, pre-blocked with 10µg/mL each anti-Siglec-5/14, anti-Siglec-6 and anti-Siglec-9, in combination with nothing (black) or 100µg/mL PSIa (dark grey) or PSV (light grey). **e**, Frequencies of serotype-specific B cells in 4 different South African combined donor PBMC pools (shades of grey). Average, standard deviation and individual measurements from two technical duplicates are shown. **f**. Pearson correlation plot showing the similarity between the technical duplicates from (e). **g**, Frequency of cells in the “empty” category (BUV615$^{+}$BUV661$^{+}$) from all donors from (Fig. 5) to indicate the background and lower limit of detection. **h**, Clusters of IgD$^{+}$IgM$^{+}$ (orange), IgD$^{+}$ (dark green), IgM$^{+}$ (light green), IgA$^{+}$ (blue) and IgG$^{+}$ (magenta) B cells on a UMAP. **i**, Heatmap showing the difference in confidence intervals for each serotype and isotype between Dutch and South African (SA) donors, assessed through a two-tailed test based on a linear regression model with wild bootstrap simulation at 9999 resamples. Bonferroni-Hochberg correction for multiple testing was employed. Stars indicate which PS-specific B cells and isotype are significantly associated to South African donors. (* $p<0.05$)

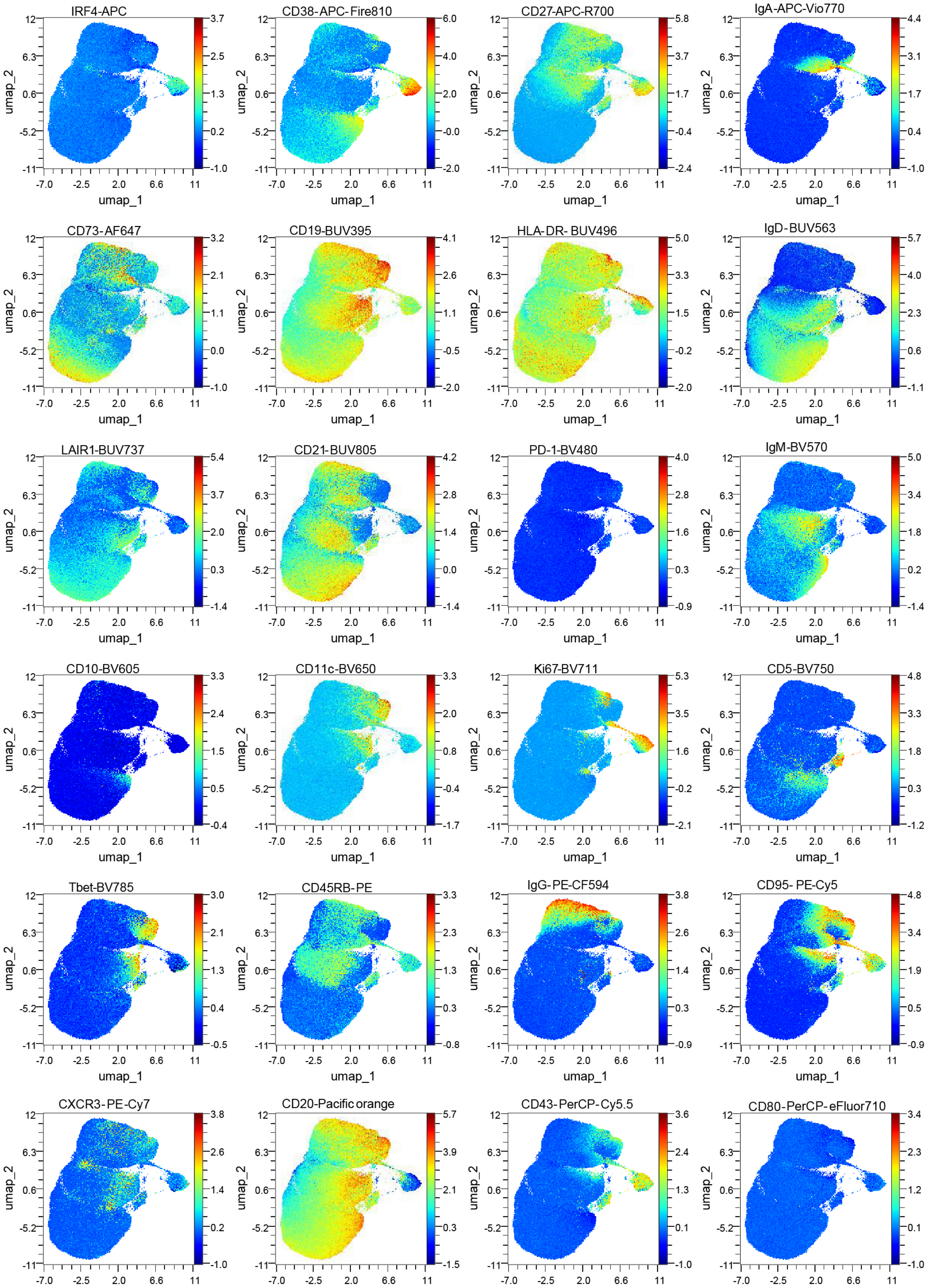
IRF4-APC
CD38-APC-Fire810
CD27-APC-R700
IgA-APC-Vio770
CD73-AF647
CD19-BUV395
HLA-DR- BUV496
IgD-BUV563
LAIR1-BUV737
CD21-BUV805
PD-1-BV480
IgM-BV570
CD10-BV605
CD11c-BV650
Ki67-BV711
CD5-BV750
Tbet-BV785
CD45RB-PE
IgG-PE-CF594
CD95- PE-Cy5
CXCR3-PE-Cy7
CD20-Pacific orange
CD43-PerCP-Cy5.5
CD80-PerCP-eFluor710
umap_1
umap_2

**Supplementary Fig. 6. Marker expression distribution across the UMAP of GBS samples.** Density plots from all clustered B cells from (Fig. 5).
