## Supplementary Tables for "Combinatorial multimer staining and spectral flow cytometry facilitate quantification and characterization of polysaccharide-specific B cell immunity"

| Supplementary table 1 | | | | | | | |
|---|---|---|---|---|---|---|---|
| **Marker/dye** | **Fluorochrome** | **Clone** | **Isotype** | **Company** | **Catalogue** | **Dilution** | **Surface/intracel.** |
| CD3 | BV510 | OKT3 | Mouse IgG2a. κ | Biolegend | 317332 | 100 | Surface |
| CD56 | BV510 | HCD56 | Mouse IgG1. κ | Biolegend | 318340 | 50 | Surface |
| CD19 | BUV395 | HIB19 | Mouse IgG1. κ | BD | 740287 | 200 | Surface |
| HLA-DR | BUV496 | G46-6 | Mouse IgG2a. κ | BD | 749866 | 200 | Surface |
| IgD | BUV563 | I-A6-2 | Mouse IgG2a. κ | BD | 741394 | 400 | Surface |
| IgM | BV570 | MHM-88 | Mouse IgG1. κ | Biolegend | 314517 | 400 | Surface |
| IgA | APC-Vio770 | IS11-8E10 | Mouse IgG1κ | Miltenyi | 130-113-999 | 1600 | Surface |
| IgG | PE-CF594 | G18-145 | Mouse IgG1. κ | BD | 562538 | 400 | Surface |
| CD138 | BUV737 | MI15 | Mouse IgG1. κ | BD | 612834 | 50 | Surface |
| CD38 | APC-Fire810 | HIT2 | Mouse IgG1. κ | Biolegend | 303550 | 100 | Surface |
| CD10 | BV605 | HI 10 a | Mouse IgG1. κ | Biolegend | 312222 | 100 | Surface |
| CD11c | BV650 | Bu15 | Mouse IgG1. κ | Biolegend | 337237 | 400 | Surface |
| CD20 | Pacific orange | HI47 | Mouse IgG3 | ThermoFisher | MHCD2030 | 20 | Surface |
| CD21 | BUV805 | BLy4 | Mouse IgG1. κ | BD | 742008 | 200 | Surface |
| CD27 | APC-R700 | M-T271 | Mouse IgG1. κ | BD | 565116 | 100 | Surface |
| CD43 | PerCP-Cy5.5 | 1G10 | Mouse IgG1. κ | BD | 563521 | 400 | Surface |
| $CD45RB_{MEM55}$ | PE | MEM-55 | Mouse IgG2b. κ | Biolegend | 310204 | 400 | Surface |
| CD5 | BV750 | L17F12 | Mouse IgG2a. κ | BD | 747090 | 200 | Surface |
| CD73 | AF647 | AD2 | Mouse IgG1 | Abcam | 243083 | 200 | Surface |
| CD95 | PE-Cy5 | DX2 | Mouse IgG1. κ | Biolegend | 305610 | 400 | Surface |
| CD80 | PerCP-eFluor710 | 16-10A1 | Hamster IgG | ThermoFisher | 46-0801-82 | 200 | Surface |
| CXCR3 | PE-Cy7 | G025H7 | Mouse IgG1. κ | Biolegend | 353719 | 1200 | Surface |
| PD-1 | BV480 | EH12.1 | Mouse IgG1. κ | BD | 566112 | 100 | Surface |
| IRF4 | APC | REA201 | Human IgG1 | Miltenyi | 130-100-915 | 200 | Intracellular |
| Caspase-3 | V450 | C92-605 | Rabbit IgG | BD | 560627 | 100 | Intracellular |
| SA-BB515 | BB515 | NA | NA | BD | 564453 | NA | NA |
| SA-BUV615 | BUV615 | NA | NA | BD | 613013 | NA | NA |
| SA-BUV661 | BUV661 | NA | NA | BD | 612979 | NA | NA |
| SA-BV421 | BV421 | NA | NA | BD | 563259 | NA | NA |
| SA-BV711 | BV711 | NA | NA | Biolegend | 405241 | NA | NA |
| SA-BV785 | BV785 | NA | NA | Biolegend | 405249 | NA | NA |
| Live/Dead | Blue | NA | NA | ThermoFisher | L34962 | 500 | NA |

**Supplementary table 1. Antibody list for Spn analysis.**

| Supplementary table 2 | | | | | | | |
|---|---|---|---|---|---|---|---|
| cluster | type | obs_estimate | unbiased_estimate | ci_limit_0.05 | ci.adj_limit_0.00357142857142857 | signif_ci | signif_ci.adj |
| elbow02 | PS3 | 0,049868808 | 0,050062022 | 0,023916796 | 0,007442109 | 1 | 1 |
| elbow03 | PS19F | 0,108454984 | 0,10828408 | 0,049139345 | 0,012581391 | 1 | 1 |
| elbow12 | PS19F | 0,028202472 | 0,028180124 | 0,015238202 | 0,006695311 | 1 | 1 |
| elbow22 | PS3 | 0,072066764 | 0,072143482 | 0,048592785 | 0,033406243 | 1 | 1 |
| elbow24 | PS3 | 0,023295111 | 0,023248339 | 0,012181427 | 0,005125825 | 1 | 1 |
| elbow24 | PS1 | 0,035488191 | 0,035559035 | 0,015258346 | 0,00296391 | 1 | 1 |
| elbow02 | PS14 | 0,063593536 | 0,063777471 | 0,008077162 | -0,027667168 | 1 | 0 |
| elbow03 | PS15B | 0,125187621 | 0,125986717 | 0,02723063 | -0,034114835 | 1 | 0 |
| elbow03 | PS7F | 0,080551301 | 0,080795597 | 0,018086978 | -0,022874351 | 1 | 0 |
| elbow03 | PS9V | 0,113897479 | 0,113303016 | 0,032383169 | -0,018956291 | 1 | 0 |
| elbow04 | PS14 | 0,116961197 | 0,116411835 | 0,012978791 | -0,05535761 | 1 | 0 |
| elbow04 | PS23F | 0,151695407 | 0,150736944 | 0,004693672 | -0,08341464 | 1 | 0 |
| elbow06 | PS6B | 0,05642263 | 0,05617144 | 0,001629459 | -0,031792753 | 1 | 0 |
| elbow07 | PS3 | 0,020844488 | 0,020567777 | 0,000304645 | -0,012431635 | 1 | 0 |
| elbow08 | PS3 | 0,027177645 | 0,02716793 | 0,00358492 | -0,010910642 | 1 | 0 |
| elbow08 | PS1 | 0,055944928 | 0,056253144 | 0,012200062 | -0,013896332 | 1 | 0 |
| elbow09 | PS3 | 0,011284919 | 0,011325884 | 0,003401641 | -0,001023324 | 1 | 0 |
| elbow11 | PS15B | 0,04261636 | 0,04277258 | 0,00278086 | -0,022983941 | 1 | 0 |
| elbow11 | PS1 | 0,041879075 | 0,042085738 | 0,015308652 | -0,001136055 | 1 | 0 |
| elbow12 | PS9V | 0,026058373 | 0,025934497 | 0,002873525 | -0,013405547 | 1 | 0 |
| elbow15 | PS6B | 0,025894977 | 0,025879619 | 0,003921525 | -0,010208267 | 1 | 0 |
| elbow16 | PS1 | 0,04441956 | 0,044269971 | 0,01507113 | -0,00394438 | 1 | 0 |
| elbow21 | PS4 | 0,16336651 | 0,163612242 | 0,055853611 | -0,012689965 | 1 | 0 |
| elbow22 | PS1 | 0,035493897 | 0,035464321 | 0,010831155 | -0,006370062 | 1 | 0 |
| elbow25 | PS6B | 0,029481139 | 0,029434947 | 0,001684514 | -0,015323105 | 1 | 0 |
| elbow27 | PS3 | 0,008515867 | 0,008528769 | 0,000366625 | -0,004924165 | 1 | 0 |
| elbow28 | PS6B | 0,027595813 | 0,027443715 | 0,001744985 | -0,014050952 | 1 | 0 |
| elbow31 | PS3 | 0,018223097 | 0,018287247 | 7,79877E-06 | -0,011587024 | 1 | 0 |
| elbow33 | PS6B | 0,025390256 | 0,025463697 | 0,000736891 | -0,016333152 | 1 | 0 |
| elbow35 | PS18C | 0,018226209 | 0,018408384 | 0,000652471 | -0,010474755 | 1 | 0 |
| elbow35 | PS6B | 0,044950402 | 0,045013448 | 0,002080034 | -0,025259934 | 1 | 0 |
| elbow01 | PS14 | 0,000901005 | 0,001147537 | -0,020697606 | -0,033079483 | 0 | 0 |
| elbow01 | PS15B | -0,007974404 | -0,008043955 | -0,019457233 | -0,027425566 | 0 | 0 |
| elbow01 | PS18C | -0,009044279 | -0,009135255 | -0,020767817 | -0,028431849 | 0 | 0 |
| elbow01 | PS19A | -0,013017286 | -0,01295709 | -0,021408072 | -0,026329184 | 0 | 0 |
| elbow01 | PS19F | 0,002932769 | 0,003084169 | -0,010630012 | -0,018191137 | 0 | 0 |
| elbow01 | PS23F | -0,020873959 | -0,020873388 | -0,028461732 | -0,033605191 | 0 | 0 |
| elbow01 | PS3 | 0,010438207 | 0,010508117 | -0,002686698 | -0,010389943 | 0 | 0 |
| elbow01 | PS4 | -0,017158296 | -0,017237891 | -0,023531908 | -0,027702311 | 0 | 0 |
| elbow01 | PS5 | 0,020334832 | 0,020306564 | -0,01645003 | -0,039669025 | 0 | 0 |
| elbow01 | PS6A | 0,028807395 | 0,02873598 | -0,00048294 | -0,018194832 | 0 | 0 |
| elbow01 | PS6B | 0,01789283 | 0,017936733 | -0,009840044 | -0,028777601 | 0 | 0 |
| elbow01 | PS7F | 0,000362533 | 0,000592495 | -0,017356755 | -0,028862615 | 0 | 0 |
| elbow01 | PS9V | -0,006490084 | -0,006521141 | -0,019913019 | -0,028332241 | 0 | 0 |
| elbow01 | PS1 | -0,007111262 | -0,007083378 | -0,016040496 | -0,022196141 | 0 | 0 |
| elbow02 | PS15B | -0,025911244 | -0,025796493 | -0,044182458 | -0,056899456 | 0 | 0 |
| elbow02 | PS18C | -0,023961212 | -0,024030492 | -0,044694655 | -0,058212191 | 0 | 0 |
| elbow02 | PS19A | 0,011036481 | 0,011207577 | -0,020718353 | -0,038857407 | 0 | 0 |
| elbow02 | PS19F | -0,036804072 | -0,036966565 | -0,054739475 | -0,067364078 | 0 | 0 |
| elbow02 | PS23F | -0,063339966 | -0,063376072 | -0,074967205 | -0,081599438 | 0 | 0 |
| elbow02 | PS4 | -0,013101118 | -0,012966148 | -0,034821354 | -0,047262649 | 0 | 0 |
| elbow02 | PS5 | -0,013989316 | -0,014100181 | -0,04468782 | -0,063453695 | 0 | 0 |
| elbow02 | PS6A | 0,063600679 | 0,063740045 | -0,005786381 | -0,049746093 | 0 | 0 |
| elbow02 | PS6B | -0,004792957 | -0,004849483 | -0,040827991 | -0,064708661 | 0 | 0 |
| elbow02 | PS7F | 0,017070649 | 0,017032449 | -0,016924516 | -0,036880843 | 0 | 0 |
| elbow02 | PS9V | -0,027182172 | -0,027131064 | -0,054073359 | -0,071568267 | 0 | 0 |
| elbow02 | PS1 | 0,003911903 | 0,003976638 | -0,021976161 | -0,03735796 | 0 | 0 |
| elbow03 | PS14 | -0,037346394 | -0,036636439 | -0,087502264 | -0,119398686 | 0 | 0 |
| elbow03 | PS18C | 0,068065697 | 0,068589569 | -0,002420526 | -0,048884909 | 0 | 0 |
| elbow03 | PS19A | 0,087481012 | 0,086767321 | -0,008344757 | -0,068786831 | 0 | 0 |
| elbow03 | PS23F | -0,206431621 | -0,206562816 | -0,234966237 | -0,2523294 | 0 | 0 |
| elbow03 | PS3 | -0,071480667 | -0,071761235 | -0,116168307 | -0,142878119 | 0 | 0 |
| elbow03 | PS4 | 0,016992575 | 0,016202454 | -0,093288933 | -0,164037279 | 0 | 0 |
| elbow03 | PS5 | -0,02505189 | -0,024883156 | -0,091102434 | -0,1379007 | 0 | 0 |
| elbow03 | PS6A | -0,026618809 | -0,026650507 | -0,081939344 | -0,116005056 | 0 | 0 |
| elbow03 | PS6B | -0,15963675 | -0,159828785 | -0,200807383 | -0,228436331 | 0 | 0 |
| elbow03 | PS1 | -0,074064538 | -0,074264844 | -0,15311025 | -0,200691639 | 0 | 0 |
| elbow04 | PS15B | 0,00048713 | 0,000488477 | -0,053122303 | -0,083367887 | 0 | 0 |

| | | | | | | | |
|---|---|---|---|---|---|---|---|
| elbow04 | PS18C | 0,038006652 | 0,038346971 | -0,025072073 | -0,059826688 | 0 | 0 |
| elbow04 | PS19A | -0,002010938 | -0,001894002 | -0,06821266 | -0,113852939 | 0 | 0 |
| elbow04 | PS19F | -0,029048317 | -0,028861634 | -0,076827498 | -0,106794692 | 0 | 0 |
| elbow04 | PS3 | -0,126517641 | -0,126401009 | -0,152805962 | -0,169394463 | 0 | 0 |
| elbow04 | PS4 | -0,04458752 | -0,045099033 | -0,13778095 | -0,193797105 | 0 | 0 |
| elbow04 | PS5 | 0,050357855 | 0,050194771 | -0,020845309 | -0,071961253 | 0 | 0 |
| elbow04 | PS6A | -0,006392878 | -0,006204533 | -0,080935996 | -0,124493725 | 0 | 0 |
| elbow04 | PS6B | -0,136216681 | -0,136385617 | -0,191760845 | -0,230345135 | 0 | 0 |
| elbow04 | PS7F | 0,082994863 | 0,083506712 | -0,029042634 | -0,101896273 | 0 | 0 |
| elbow04 | PS9V | 0,015536263 | 0,016031396 | -0,039673278 | -0,073288816 | 0 | 0 |
| elbow04 | PS1 | -0,111265392 | -0,111556424 | -0,165593866 | -0,203649374 | 0 | 0 |
| elbow05 | PS14 | -0,001477383 | -0,001440723 | -0,005913477 | -0,008480889 | 0 | 0 |
| elbow05 | PS15B | 0,005521711 | 0,005501013 | -0,001986589 | -0,006363639 | 0 | 0 |
| elbow05 | PS18C | 0,022279066 | 0,0222205 | -0,012994016 | -0,033552769 | 0 | 0 |
| elbow05 | PS19A | -0,001710483 | -0,001726122 | -0,005862331 | -0,008539453 | 0 | 0 |
| elbow05 | PS19F | 0,002514459 | 0,002553835 | -0,004326887 | -0,008973913 | 0 | 0 |
| elbow05 | PS23F | -0,004041485 | -0,00403857 | -0,00746755 | -0,009601222 | 0 | 0 |
| elbow05 | PS3 | -0,002225246 | -0,002209179 | -0,006307889 | -0,008854296 | 0 | 0 |
| elbow05 | PS4 | -0,004041485 | -0,004059163 | -0,007419129 | -0,009501869 | 0 | 0 |
| elbow05 | PS5 | -0,004041485 | -0,004023638 | -0,00749648 | -0,009747717 | 0 | 0 |
| elbow05 | PS6A | -0,004041485 | -0,004024713 | -0,007472986 | -0,009449171 | 0 | 0 |
| elbow05 | PS6B | -0,004041485 | -0,004028936 | -0,007460708 | -0,009488397 | 0 | 0 |
| elbow05 | PS7F | -0,003480003 | -0,003521749 | -0,006747566 | -0,008792659 | 0 | 0 |
| elbow05 | PS9V | -0,001388965 | -0,001407292 | -0,006822191 | -0,01042774 | 0 | 0 |
| elbow05 | PS1 | 0,000174269 | 0,000212323 | -0,004927899 | -0,008037252 | 0 | 0 |
| elbow06 | PS14 | 0,005611089 | 0,005748865 | -0,018375215 | -0,033182685 | 0 | 0 |
| elbow06 | PS15B | 0,013983793 | 0,014018852 | -0,011278689 | -0,027584296 | 0 | 0 |
| elbow06 | PS18C | 0,002523045 | 0,002444873 | -0,017613374 | -0,031213071 | 0 | 0 |
| elbow06 | PS19A | -0,009331098 | -0,009400145 | -0,029648596 | -0,042491577 | 0 | 0 |
| elbow06 | PS19F | -0,004019559 | -0,004246017 | -0,020845567 | -0,032496299 | 0 | 0 |
| elbow06 | PS23F | -0,035056191 | -0,035091967 | -0,043313222 | -0,048900279 | 0 | 0 |
| elbow06 | PS3 | -0,012284875 | -0,012301109 | -0,025665342 | -0,034690275 | 0 | 0 |
| elbow06 | PS4 | 0,010395168 | 0,010361932 | -0,015371496 | -0,031263065 | 0 | 0 |
| elbow06 | PS5 | 0,041766986 | 0,041510363 | -0,018638287 | -0,05458044 | 0 | 0 |
| elbow06 | PS6A | -0,016581743 | -0,016527269 | -0,034662192 | -0,047731774 | 0 | 0 |
| elbow06 | PS7F | -0,023556288 | -0,023596789 | -0,033434701 | -0,03960302 | 0 | 0 |
| elbow06 | PS9V | -0,010164704 | -0,010156059 | -0,022951646 | -0,031196173 | 0 | 0 |
| elbow06 | PS1 | -0,019708255 | -0,019748093 | -0,033505203 | -0,042902762 | 0 | 0 |
| elbow07 | PS14 | 0,00256465 | 0,002336404 | -0,025794984 | -0,043169677 | 0 | 0 |
| elbow07 | PS15B | -0,008545024 | -0,008677797 | -0,026016297 | -0,037352804 | 0 | 0 |
| elbow07 | PS18C | -0,018138043 | -0,018244491 | -0,032821939 | -0,041349064 | 0 | 0 |
| elbow07 | PS19A | 0,007204499 | 0,007556201 | -0,018545119 | -0,034496209 | 0 | 0 |
| elbow07 | PS19F | 0,012762532 | 0,012715004 | -0,006325608 | -0,018000211 | 0 | 0 |
| elbow07 | PS23F | -0,018994496 | -0,018709491 | -0,050552217 | -0,070528604 | 0 | 0 |
| elbow07 | PS4 | 0,030982123 | 0,030890567 | -0,016698841 | -0,049382337 | 0 | 0 |
| elbow07 | PS5 | 0,002101075 | 0,00244554 | -0,035639975 | -0,058468552 | 0 | 0 |
| elbow07 | PS6A | -0,018445045 | -0,018366365 | -0,039863871 | -0,05334391 | 0 | 0 |
| elbow07 | PS6B | -0,032308105 | -0,032262012 | -0,046712747 | -0,056533857 | 0 | 0 |
| elbow07 | PS7F | 0,01245204 | 0,012474625 | -0,022414939 | -0,043940004 | 0 | 0 |
| elbow07 | PS9V | 0,007973694 | 0,008095349 | -0,007443563 | -0,017564322 | 0 | 0 |
| elbow07 | PS1 | -0,000454387 | -0,000520707 | -0,01885771 | -0,030496452 | 0 | 0 |
| elbow08 | PS14 | 0,018376802 | 0,018447093 | -0,006176979 | -0,021856423 | 0 | 0 |
| elbow08 | PS15B | -0,008298829 | -0,008293921 | -0,025386867 | -0,036778548 | 0 | 0 |
| elbow08 | PS18C | -0,002447435 | -0,002457699 | -0,019798989 | -0,031026903 | 0 | 0 |
| elbow08 | PS19A | 0,009140999 | 0,009200356 | -0,011250487 | -0,024074735 | 0 | 0 |
| elbow08 | PS19F | -0,029891108 | -0,0299162 | -0,038823994 | -0,043889401 | 0 | 0 |
| elbow08 | PS23F | -0,022961038 | -0,022881697 | -0,037602134 | -0,047252463 | 0 | 0 |
| elbow08 | PS4 | 0,006428624 | 0,0063339 | -0,027814294 | -0,047929876 | 0 | 0 |
| elbow08 | PS5 | -0,021462536 | -0,021449471 | -0,036660952 | -0,04679275 | 0 | 0 |
| elbow08 | PS6A | 0,001927655 | 0,001871657 | -0,015057971 | -0,02567728 | 0 | 0 |
| elbow08 | PS6B | 0,009517595 | 0,009678141 | -0,020617023 | -0,0402241 | 0 | 0 |
| elbow08 | PS7F | -0,024267933 | -0,024140826 | -0,032419047 | -0,037158473 | 0 | 0 |
| elbow08 | PS9V | -0,01918537 | -0,019154263 | -0,032857074 | -0,041496432 | 0 | 0 |
| elbow09 | PS14 | 0,000151337 | 0,000172221 | -0,00853391 | -0,013847202 | 0 | 0 |
| elbow09 | PS15B | -0,005309468 | -0,005324935 | -0,010927894 | -0,014299956 | 0 | 0 |
| elbow09 | PS18C | 0,00137068 | 0,001374834 | -0,007145024 | -0,012542002 | 0 | 0 |
| elbow09 | PS19A | -0,002708298 | -0,002758156 | -0,010089207 | -0,014420954 | 0 | 0 |
| elbow09 | PS19F | 0,003069679 | 0,003061351 | -0,007407522 | -0,013422628 | 0 | 0 |
| elbow09 | PS23F | -0,008805972 | -0,008840488 | -0,012416569 | -0,014890869 | 0 | 0 |
| elbow09 | PS4 | -0,004647967 | -0,004710162 | -0,011323807 | -0,015569353 | 0 | 0 |

| | | | | | | | |
|---|---|---|---|---|---|---|---|
| elbow09 | PS5 | -0,008805972 | -0,008796998 | -0,012556398 | -0,015029666 | 0 | 0 |
| elbow09 | PS6A | 0,010269824 | 0,010333657 | -0,007938609 | -0,019776513 | 0 | 0 |
| elbow09 | PS6B | 0,009356422 | 0,009272434 | -0,007607565 | -0,018008727 | 0 | 0 |
| elbow09 | PS7F | -0,001098977 | -0,001144593 | -0,006966958 | -0,011163311 | 0 | 0 |
| elbow09 | PS9V | -0,006641925 | -0,006612058 | -0,010213339 | -0,012369475 | 0 | 0 |
| elbow09 | PS1 | 0,002515718 | 0,002534956 | -0,005821518 | -0,010926338 | 0 | 0 |
| elbow10 | PS14 | -0,009361596 | -0,009487626 | -0,02132529 | -0,028704085 | 0 | 0 |
| elbow10 | PS15B | -0,003761982 | -0,003731003 | -0,016634688 | -0,024609961 | 0 | 0 |
| elbow10 | PS18C | -0,006843811 | -0,006843321 | -0,021126164 | -0,02990647 | 0 | 0 |
| elbow10 | PS19A | 0,001301959 | 0,001299415 | -0,013378498 | -0,022780421 | 0 | 0 |
| elbow10 | PS19F | -0,01377855 | -0,013923292 | -0,024469825 | -0,0309873 | 0 | 0 |
| elbow10 | PS23F | 0,079940205 | 0,081254009 | -0,034504217 | -0,103257152 | 0 | 0 |
| elbow10 | PS3 | -0,008518819 | -0,008590559 | -0,019919451 | -0,02761138 | 0 | 0 |
| elbow10 | PS4 | 0,009427384 | 0,009170085 | -0,028164792 | -0,05215652 | 0 | 0 |
| elbow10 | PS5 | -0,016213641 | -0,016183814 | -0,026693662 | -0,03317421 | 0 | 0 |
| elbow10 | PS6A | -0,000691663 | -0,000751084 | -0,017768321 | -0,028907827 | 0 | 0 |
| elbow10 | PS6B | -0,016213641 | -0,016162776 | -0,02699914 | -0,034136654 | 0 | 0 |
| elbow10 | PS7F | 0,006486719 | 0,006594672 | -0,017339479 | -0,033014461 | 0 | 0 |
| elbow10 | PS9V | -0,009102368 | -0,009109184 | -0,020488525 | -0,02774546 | 0 | 0 |
| elbow10 | PS1 | -0,012670194 | -0,012534106 | -0,024844651 | -0,03242655 | 0 | 0 |
| elbow11 | PS14 | -0,006854511 | -0,006882172 | -0,030890895 | -0,044859074 | 0 | 0 |
| elbow11 | PS18C | -0,018886157 | -0,01865799 | -0,04391423 | -0,060304578 | 0 | 0 |
| elbow11 | PS19A | -0,000847944 | -0,001221645 | -0,02875579 | -0,047132181 | 0 | 0 |
| elbow11 | PS19F | -0,038561046 | -0,038432561 | -0,050796794 | -0,058044868 | 0 | 0 |
| elbow11 | PS23F | -0,023580954 | -0,023644936 | -0,0768755 | -0,107624418 | 0 | 0 |
| elbow11 | PS3 | -0,000817817 | -0,000774354 | -0,022429583 | -0,036705795 | 0 | 0 |
| elbow11 | PS4 | -0,031776547 | -0,031757137 | -0,061983056 | -0,080000097 | 0 | 0 |
| elbow11 | PS5 | 0,045849616 | 0,045582129 | -0,035085422 | -0,086618851 | 0 | 0 |
| elbow11 | PS6A | -0,011758856 | -0,01196638 | -0,036030184 | -0,05219964 | 0 | 0 |
| elbow11 | PS6B | 0,030077354 | 0,030319628 | -0,019880138 | -0,050081974 | 0 | 0 |
| elbow11 | PS7F | -0,007431324 | -0,007548399 | -0,035499019 | -0,053468884 | 0 | 0 |
| elbow11 | PS9V | -0,019907248 | -0,019786164 | -0,039732102 | -0,052895026 | 0 | 0 |
| elbow12 | PS14 | -0,002779618 | -0,002703346 | -0,01150837 | -0,017020662 | 0 | 0 |
| elbow12 | PS15B | -0,003830046 | -0,003820975 | -0,011078451 | -0,015710512 | 0 | 0 |
| elbow12 | PS18C | -0,00261595 | -0,002714293 | -0,011163151 | -0,016530844 | 0 | 0 |
| elbow12 | PS19A | 0,023663295 | 0,023554379 | -0,005199781 | -0,023326121 | 0 | 0 |
| elbow12 | PS23F | -0,014850179 | -0,014876874 | -0,018750452 | -0,021122258 | 0 | 0 |
| elbow12 | PS3 | -0,00593752 | -0,005931101 | -0,011393648 | -0,014791078 | 0 | 0 |
| elbow12 | PS4 | -0,009055514 | -0,009120163 | -0,016469102 | -0,021032805 | 0 | 0 |
| elbow12 | PS5 | -0,011353676 | -0,011380547 | -0,017333012 | -0,020963308 | 0 | 0 |
| elbow12 | PS6A | -0,012652377 | -0,012650104 | -0,017454407 | -0,020131996 | 0 | 0 |
| elbow12 | PS6B | -0,014850179 | -0,014844616 | -0,018840988 | -0,021350063 | 0 | 0 |
| elbow12 | PS7F | -0,00068511 | -0,000627267 | -0,012226886 | -0,019796272 | 0 | 0 |
| elbow12 | PS1 | 0,00068603 | 0,000698561 | -0,014897686 | -0,024532649 | 0 | 0 |
| elbow13 | PS14 | -0,00175778 | -0,001747904 | -0,003321999 | -0,004298834 | 0 | 0 |
| elbow13 | PS15B | -0,00175778 | -0,001768157 | -0,003320728 | -0,004283467 | 0 | 0 |
| elbow13 | PS18C | -0,00175778 | -0,001749278 | -0,003288523 | -0,004344233 | 0 | 0 |
| elbow13 | PS19A | 0,002052512 | 0,002058438 | -0,001398786 | -0,003466896 | 0 | 0 |
| elbow13 | PS19F | 0,002312391 | 0,002309545 | -0,002322506 | -0,00514014 | 0 | 0 |
| elbow13 | PS23F | -0,00175778 | -0,0017425 | -0,003307637 | -0,004236588 | 0 | 0 |
| elbow13 | PS3 | -0,000166268 | -0,000182737 | -0,002503025 | -0,004002567 | 0 | 0 |
| elbow13 | PS4 | -0,00175778 | -0,001748989 | -0,003338993 | -0,004251198 | 0 | 0 |
| elbow13 | PS5 | 0,005235227 | 0,005311777 | -0,00463698 | -0,0107617 | 0 | 0 |
| elbow13 | PS6A | -0,00175778 | -0,001750555 | -0,003278217 | -0,004184296 | 0 | 0 |
| elbow13 | PS6B | -0,00175778 | -0,001762408 | -0,003309494 | -0,004359399 | 0 | 0 |
| elbow13 | PS7F | 0,002088374 | 0,002096209 | -0,003163468 | -0,006523294 | 0 | 0 |
| elbow13 | PS9V | 0,001738723 | 0,001726296 | -0,003126744 | -0,006278752 | 0 | 0 |
| elbow13 | PS1 | -0,000956498 | -0,00094533 | -0,002576545 | -0,003564522 | 0 | 0 |
| elbow14 | PS14 | -0,003288852 | -0,003255458 | -0,010625022 | -0,01513435 | 0 | 0 |
| elbow14 | PS15B | 0,004783396 | 0,004770527 | -0,004129166 | -0,009536757 | 0 | 0 |
| elbow14 | PS18C | -0,001216722 | -0,001271748 | -0,009568571 | -0,014849381 | 0 | 0 |
| elbow14 | PS19A | -0,009687418 | -0,009688945 | -0,014245534 | -0,017012767 | 0 | 0 |
| elbow14 | PS19F | 0,009122758 | 0,008971318 | -0,006043822 | -0,015978295 | 0 | 0 |
| elbow14 | PS23F | 0,035184377 | 0,034992045 | -0,010051369 | -0,038008125 | 0 | 0 |
| elbow14 | PS3 | -0,004251077 | -0,004234026 | -0,009728344 | -0,013573736 | 0 | 0 |
| elbow14 | PS4 | -0,009687418 | -0,00963852 | -0,014354522 | -0,017347563 | 0 | 0 |
| elbow14 | PS5 | -0,009687418 | -0,009630542 | -0,01430219 | -0,017379418 | 0 | 0 |
| elbow14 | PS6A | -0,002843474 | -0,002907863 | -0,012635129 | -0,018902469 | 0 | 0 |
| elbow14 | PS6B | 0,012290604 | 0,012321913 | -0,008860331 | -0,021951512 | 0 | 0 |
| elbow14 | PS7F | -0,004751253 | -0,004753332 | -0,010185935 | -0,013538837 | 0 | 0 |

| | | | | | | | |
|---|---|---|---|---|---|---|---|
| elbow14 | PS9V | -0,006849888 | -0,006901111 | -0,012353016 | -0,015840879 | 0 | 0 |
| elbow14 | PS1 | -0,009117617 | -0,009111463 | -0,013938517 | -0,017084962 | 0 | 0 |
| elbow15 | PS14 | -0,003256427 | -0,003251055 | -0,005887417 | -0,007575313 | 0 | 0 |
| elbow15 | PS15B | -0,003256427 | -0,003253842 | -0,005847828 | -0,007356347 | 0 | 0 |
| elbow15 | PS18C | 0,004032655 | 0,004106689 | -0,002471194 | -0,006281463 | 0 | 0 |
| elbow15 | PS19A | -0,003256427 | -0,003256944 | -0,005756347 | -0,007472467 | 0 | 0 |
| elbow15 | PS19F | -0,003256427 | -0,003269656 | -0,005816866 | -0,007350582 | 0 | 0 |
| elbow15 | PS23F | -0,003256427 | -0,003259623 | -0,005784575 | -0,007424552 | 0 | 0 |
| elbow15 | PS3 | -0,003256427 | -0,003247818 | -0,005775373 | -0,007439209 | 0 | 0 |
| elbow15 | PS4 | 0,002238079 | 0,002214969 | -0,006123456 | -0,01116145 | 0 | 0 |
| elbow15 | PS5 | -0,003256427 | -0,003275209 | -0,005814006 | -0,007485228 | 0 | 0 |
| elbow15 | PS6A | -0,003256427 | -0,003239279 | -0,005816625 | -0,007465753 | 0 | 0 |
| elbow15 | PS7F | 4,07774E-05 | 4,69903E-05 | -0,002838266 | -0,0045944 | 0 | 0 |
| elbow15 | PS9V | -0,002898645 | -0,002900932 | -0,005578299 | -0,007255707 | 0 | 0 |
| elbow15 | PS1 | -0,003256427 | -0,003271585 | -0,005790893 | -0,007530236 | 0 | 0 |
| elbow16 | PS14 | -0,00954428 | -0,009369101 | -0,025555227 | -0,035640711 | 0 | 0 |
| elbow16 | PS15B | 0,021009479 | 0,021132416 | -0,015347907 | -0,041005652 | 0 | 0 |
| elbow16 | PS18C | -0,007552277 | -0,007745923 | -0,030323023 | -0,044433098 | 0 | 0 |
| elbow16 | PS19A | -0,003263716 | -0,00315907 | -0,023770259 | -0,037698209 | 0 | 0 |
| elbow16 | PS19F | -0,012712505 | -0,012653929 | -0,034113907 | -0,046991346 | 0 | 0 |
| elbow16 | PS23F | -0,036706175 | -0,03678083 | -0,045628278 | -0,052184823 | 0 | 0 |
| elbow16 | PS3 | 0,016050549 | 0,016214196 | -0,011370389 | -0,028108501 | 0 | 0 |
| elbow16 | PS4 | -0,020084366 | -0,020229649 | -0,036429422 | -0,046545718 | 0 | 0 |
| elbow16 | PS5 | 0,014109676 | 0,013955221 | -0,025747567 | -0,052535307 | 0 | 0 |
| elbow16 | PS6A | -0,008482036 | -0,008400738 | -0,03088539 | -0,046641267 | 0 | 0 |
| elbow16 | PS6B | -0,018543782 | -0,018567232 | -0,038524848 | -0,050840003 | 0 | 0 |
| elbow16 | PS7F | -0,003263435 | -0,003048878 | -0,028135776 | -0,04399703 | 0 | 0 |
| elbow16 | PS9V | 0,024563307 | 0,02438194 | -0,00799473 | -0,029404895 | 0 | 0 |
| elbow17 | PS14 | -0,003023774 | -0,003018643 | -0,006304901 | -0,008578544 | 0 | 0 |
| elbow17 | PS15B | -0,003023774 | -0,00301841 | -0,006230889 | -0,008265925 | 0 | 0 |
| elbow17 | PS18C | -0,001515478 | -0,001504076 | -0,005554026 | -0,008471054 | 0 | 0 |
| elbow17 | PS19A | -0,003023774 | -0,003042784 | -0,006229407 | -0,008141231 | 0 | 0 |
| elbow17 | PS19F | -0,003023774 | -0,003052441 | -0,006246368 | -0,008538263 | 0 | 0 |
| elbow17 | PS23F | -0,003023774 | -0,002968454 | -0,006361797 | -0,008318056 | 0 | 0 |
| elbow17 | PS3 | -0,000660777 | -0,000635547 | -0,004992675 | -0,007560891 | 0 | 0 |
| elbow17 | PS4 | -0,003023774 | -0,003023199 | -0,0062566 | -0,008296199 | 0 | 0 |
| elbow17 | PS5 | -0,003023774 | -0,003018706 | -0,006289608 | -0,008264059 | 0 | 0 |
| elbow17 | PS6A | -0,003023774 | -0,003028651 | -0,006189991 | -0,008362269 | 0 | 0 |
| elbow17 | PS6B | 0,035437765 | 0,035558824 | -0,004012777 | -0,028679834 | 0 | 0 |
| elbow17 | PS7F | -0,003023774 | -0,00304164 | -0,006309884 | -0,008576465 | 0 | 0 |
| elbow17 | PS9V | -0,003023774 | -0,003015057 | -0,006296336 | -0,008319487 | 0 | 0 |
| elbow17 | PS1 | -0,003023774 | -0,003050424 | -0,00621367 | -0,008237676 | 0 | 0 |
| elbow18 | PS14 | -0,002930403 | -0,002952718 | -0,00558016 | -0,00726331 | 0 | 0 |
| elbow18 | PS15B | -0,002930403 | -0,002943756 | -0,005548062 | -0,007215396 | 0 | 0 |
| elbow18 | PS18C | -0,002930403 | -0,002947825 | -0,005553337 | -0,007157587 | 0 | 0 |
| elbow18 | PS19A | -0,002930403 | -0,002914541 | -0,005568667 | -0,00726701 | 0 | 0 |
| elbow18 | PS19F | -0,002930403 | -0,002945216 | -0,005531389 | -0,007088846 | 0 | 0 |
| elbow18 | PS23F | -0,002930403 | -0,002940449 | -0,005549725 | -0,007360436 | 0 | 0 |
| elbow18 | PS3 | -0,002930403 | -0,002936979 | -0,00556406 | -0,007322725 | 0 | 0 |
| elbow18 | PS4 | -0,002930403 | -0,002918885 | -0,00560995 | -0,007184191 | 0 | 0 |
| elbow18 | PS5 | -0,002930403 | -0,002930581 | -0,005585521 | -0,007247068 | 0 | 0 |
| elbow18 | PS6A | -0,002930403 | -0,002936512 | -0,005527392 | -0,007117934 | 0 | 0 |
| elbow18 | PS6B | 0,025274725 | 0,025507119 | -0,003024787 | -0,020293257 | 0 | 0 |
| elbow18 | PS7F | 0,00989011 | 0,009870173 | -0,008592397 | -0,020025634 | 0 | 0 |
| elbow18 | PS9V | -0,002930403 | -0,002929066 | -0,005548601 | -0,00722774 | 0 | 0 |
| elbow18 | PS1 | -0,002930403 | -0,002924189 | -0,005645755 | -0,007335092 | 0 | 0 |
| elbow19 | PS14 | -0,004151702 | -0,004131676 | -0,00736259 | -0,009460495 | 0 | 0 |
| elbow19 | PS15B | -0,004151702 | -0,004173828 | -0,007322739 | -0,009239147 | 0 | 0 |
| elbow19 | PS18C | -0,004151702 | -0,004169313 | -0,007311185 | -0,009236499 | 0 | 0 |
| elbow19 | PS19A | -0,004151702 | -0,00412915 | -0,007295264 | -0,009435459 | 0 | 0 |
| elbow19 | PS19F | 0,006809836 | 0,006793806 | -0,004722778 | -0,012499817 | 0 | 0 |
| elbow19 | PS23F | 0,026068078 | 0,025904628 | -0,004264107 | -0,023501063 | 0 | 0 |
| elbow19 | PS3 | -0,00388645 | -0,003893237 | -0,007199353 | -0,009121959 | 0 | 0 |
| elbow19 | PS4 | -0,0020727 | -0,002005732 | -0,005857186 | -0,008295413 | 0 | 0 |
| elbow19 | PS5 | -0,004151702 | -0,004157902 | -0,007260037 | -0,0092628 | 0 | 0 |
| elbow19 | PS6A | -0,001193122 | -0,001217732 | -0,005534683 | -0,008267942 | 0 | 0 |
| elbow19 | PS6B | 0,005463683 | 0,005392239 | -0,008184779 | -0,015891906 | 0 | 0 |
| elbow19 | PS7F | -0,002127411 | -0,002085489 | -0,006512217 | -0,009217818 | 0 | 0 |
| elbow19 | PS9V | -0,004151702 | -0,004178683 | -0,007286981 | -0,009255462 | 0 | 0 |
| elbow19 | PS1 | -0,004151702 | -0,004135207 | -0,007291948 | -0,009416417 | 0 | 0 |

| | | | | | | | |
|---|---|---|---|---|---|---|---|
| elbow20 | PS14 | -0,008026028 | -0,007989885 | -0,012432673 | -0,015000715 | 0 | 0 |
| elbow20 | PS15B | -0,008026028 | -0,008057829 | -0,012291736 | -0,015209653 | 0 | 0 |
| elbow20 | PS18C | -0,008026028 | -0,007994137 | -0,012400241 | -0,015094252 | 0 | 0 |
| elbow20 | PS19A | -0,002108868 | -0,002036176 | -0,011409095 | -0,016889196 | 0 | 0 |
| elbow20 | PS19F | 0,018267678 | 0,0181284 | -0,002389273 | -0,01421565 | 0 | 0 |
| elbow20 | PS23F | -0,008026028 | -0,008027054 | -0,012344557 | -0,014990091 | 0 | 0 |
| elbow20 | PS3 | 0,001588776 | 0,001546856 | -0,008499136 | -0,014693195 | 0 | 0 |
| elbow20 | PS4 | 0,005959986 | 0,005973713 | -0,015047765 | -0,029238029 | 0 | 0 |
| elbow20 | PS5 | 0,011204741 | 0,011123628 | -0,016646692 | -0,0339571 | 0 | 0 |
| elbow20 | PS6A | -0,00033372 | -0,000401969 | -0,012807531 | -0,020324399 | 0 | 0 |
| elbow20 | PS6B | -0,008026028 | -0,008004639 | -0,012345956 | -0,014896915 | 0 | 0 |
| elbow20 | PS7F | -0,008026028 | -0,00802845 | -0,012360752 | -0,015221905 | 0 | 0 |
| elbow20 | PS9V | 0,007358588 | 0,007439117 | -0,015043559 | -0,029304237 | 0 | 0 |
| elbow20 | PS1 | 0,006218986 | 0,006259746 | -0,006810191 | -0,014238123 | 0 | 0 |
| elbow21 | PS14 | -0,013999108 | -0,014025309 | -0,023359877 | -0,0292993 | 0 | 0 |
| elbow21 | PS15B | -0,013999108 | -0,013930761 | -0,023498595 | -0,029734499 | 0 | 0 |
| elbow21 | PS18C | -0,013999108 | -0,013999123 | -0,023412349 | -0,029398662 | 0 | 0 |
| elbow21 | PS19A | -0,005257849 | -0,005293384 | -0,018081632 | -0,026721076 | 0 | 0 |
| elbow21 | PS19F | -0,013999108 | -0,013915179 | -0,023465261 | -0,029713524 | 0 | 0 |
| elbow21 | PS23F | -0,013999108 | -0,014014978 | -0,023335272 | -0,029565806 | 0 | 0 |
| elbow21 | PS3 | -0,013733856 | -0,013690455 | -0,023336512 | -0,029469908 | 0 | 0 |
| elbow21 | PS5 | -0,013999108 | -0,013917095 | -0,023717188 | -0,029693586 | 0 | 0 |
| elbow21 | PS6A | -0,004383723 | -0,004405558 | -0,020230029 | -0,030693921 | 0 | 0 |
| elbow21 | PS6B | -0,013999108 | -0,013982328 | -0,023359162 | -0,029053226 | 0 | 0 |
| elbow21 | PS7F | -0,013999108 | -0,014010714 | -0,023535246 | -0,029590195 | 0 | 0 |
| elbow21 | PS9V | -0,013999108 | -0,01399414 | -0,023294269 | -0,029211048 | 0 | 0 |
| elbow21 | PS1 | -0,013999108 | -0,013980569 | -0,023321075 | -0,029516421 | 0 | 0 |
| elbow22 | PS14 | -0,000111179 | -8,49378E-05 | -0,018401301 | -0,030708997 | 0 | 0 |
| elbow22 | PS15B | 0,00023489 | 0,000116467 | -0,013964764 | -0,022765513 | 0 | 0 |
| elbow22 | PS18C | -0,020137894 | -0,020214731 | -0,031331259 | -0,03895598 | 0 | 0 |
| elbow22 | PS19A | -0,001227934 | -0,001178531 | -0,0172412 | -0,026996362 | 0 | 0 |
| elbow22 | PS19F | 0,003498027 | 0,00329387 | -0,016577539 | -0,027835991 | 0 | 0 |
| elbow22 | PS23F | -0,029779127 | -0,029847923 | -0,037707332 | -0,042468133 | 0 | 0 |
| elbow22 | PS4 | -0,019512797 | -0,019488795 | -0,0313586 | -0,038686309 | 0 | 0 |
| elbow22 | PS5 | -0,000641598 | -0,000441867 | -0,034420612 | -0,05583635 | 0 | 0 |
| elbow22 | PS6A | 0,000198094 | 0,000220451 | -0,019218642 | -0,031952905 | 0 | 0 |
| elbow22 | PS6B | -0,029779127 | -0,02974449 | -0,037842139 | -0,043011834 | 0 | 0 |
| elbow22 | PS7F | -0,006909877 | -0,006826517 | -0,028847614 | -0,043170364 | 0 | 0 |
| elbow22 | PS9V | -0,00339214 | -0,003280348 | -0,016578008 | -0,02520126 | 0 | 0 |
| elbow23 | PS14 | -0,004244454 | -0,004249947 | -0,006683476 | -0,008167148 | 0 | 0 |
| elbow23 | PS15B | -0,004244454 | -0,004257156 | -0,006690627 | -0,008260811 | 0 | 0 |
| elbow23 | PS18C | -0,004244454 | -0,004258881 | -0,006747239 | -0,008294046 | 0 | 0 |
| elbow23 | PS19A | -0,004244454 | -0,004232894 | -0,00668492 | -0,008328068 | 0 | 0 |
| elbow23 | PS19F | -0,004244454 | -0,004220795 | -0,006696249 | -0,008239953 | 0 | 0 |
| elbow23 | PS23F | 0,006744557 | 0,006623276 | -0,00846677 | -0,018622734 | 0 | 0 |
| elbow23 | PS3 | -0,000786979 | -0,000799786 | -0,004434306 | -0,006819665 | 0 | 0 |
| elbow23 | PS4 | -0,004244454 | -0,004233273 | -0,006741563 | -0,008250908 | 0 | 0 |
| elbow23 | PS5 | -0,002412952 | -0,002386303 | -0,006102172 | -0,008554439 | 0 | 0 |
| elbow23 | PS6A | 0,00533571 | 0,005246771 | -0,00485122 | -0,011307488 | 0 | 0 |
| elbow23 | PS6B | 0,014986316 | 0,014867894 | -0,003473834 | -0,014615076 | 0 | 0 |
| elbow23 | PS7F | -0,004244454 | -0,00428102 | -0,006687271 | -0,008253217 | 0 | 0 |
| elbow23 | PS9V | -0,001591934 | -0,001557535 | -0,006277316 | -0,009382479 | 0 | 0 |
| elbow23 | PS1 | 0,007436458 | 0,007506192 | -0,003090762 | -0,010136451 | 0 | 0 |
| elbow24 | PS14 | -0,003864471 | -0,003821904 | -0,018184046 | -0,027842056 | 0 | 0 |
| elbow24 | PS15B | -0,009370744 | -0,009426278 | -0,01761936 | -0,022906963 | 0 | 0 |
| elbow24 | PS18C | -0,003576484 | -0,003723076 | -0,013167943 | -0,019708968 | 0 | 0 |
| elbow24 | PS19A | -0,006416296 | -0,006245617 | -0,017158904 | -0,024554419 | 0 | 0 |
| elbow24 | PS19F | 0,004132041 | 0,00411681 | -0,009269322 | -0,017680425 | 0 | 0 |
| elbow24 | PS23F | -0,005258727 | -0,005289069 | -0,020153488 | -0,029641791 | 0 | 0 |
| elbow24 | PS4 | -0,012584734 | -0,01256084 | -0,018069988 | -0,02132117 | 0 | 0 |
| elbow24 | PS5 | -0,009088231 | -0,009039002 | -0,017750839 | -0,023294639 | 0 | 0 |
| elbow24 | PS6A | -0,00855543 | -0,008444108 | -0,021331236 | -0,029801877 | 0 | 0 |
| elbow24 | PS6B | 0,005119783 | 0,005159047 | -0,016689772 | -0,030376699 | 0 | 0 |
| elbow24 | PS7F | -0,000322686 | -0,000304965 | -0,019146475 | -0,03106339 | 0 | 0 |
| elbow24 | PS9V | -0,008997323 | -0,009013463 | -0,016472192 | -0,021381294 | 0 | 0 |
| elbow25 | PS14 | -0,011356656 | -0,011396531 | -0,01800395 | -0,022333164 | 0 | 0 |
| elbow25 | PS15B | -0,011356656 | -0,011352393 | -0,01809577 | -0,022206034 | 0 | 0 |
| elbow25 | PS18C | -0,007693652 | -0,007689719 | -0,015972279 | -0,021501061 | 0 | 0 |
| elbow25 | PS19A | -0,011356656 | -0,011408787 | -0,017968706 | -0,022164608 | 0 | 0 |
| elbow25 | PS19F | 0,020996076 | 0,02075912 | -0,019839956 | -0,043851625 | 0 | 0 |

| | | | | | | | |
|---|---|---|---|---|---|---|---|
| elbow25 | PS23F | 0,027104883 | 0,027902876 | -0,030026592 | -0,068269603 | 0 | 0 |
| elbow25 | PS3 | -0,003630829 | -0,003582568 | -0,011496878 | -0,016206662 | 0 | 0 |
| elbow25 | PS4 | -0,004363649 | -0,004303971 | -0,017160345 | -0,025335443 | 0 | 0 |
| elbow25 | PS5 | -0,011356656 | -0,011399139 | -0,017972269 | -0,021964222 | 0 | 0 |
| elbow25 | PS6A | 0,011720267 | 0,011646645 | -0,022904185 | -0,045295662 | 0 | 0 |
| elbow25 | PS7F | -0,010602508 | -0,01054587 | -0,017430535 | -0,021655276 | 0 | 0 |
| elbow25 | PS9V | -0,011356656 | -0,0112598 | -0,018007638 | -0,022122451 | 0 | 0 |
| elbow25 | PS1 | -0,00622845 | -0,006192538 | -0,014794541 | -0,019755576 | 0 | 0 |
| elbow26 | PS14 | -0,017555353 | -0,017561674 | -0,027892228 | -0,034749956 | 0 | 0 |
| elbow26 | PS15B | -0,017555353 | -0,017599142 | -0,027985998 | -0,034536769 | 0 | 0 |
| elbow26 | PS18C | -0,010549076 | -0,010498776 | -0,023010415 | -0,031152828 | 0 | 0 |
| elbow26 | PS19A | -0,017555353 | -0,01747356 | -0,028075894 | -0,034576961 | 0 | 0 |
| elbow26 | PS19F | -0,0044085 | -0,004381807 | -0,019925434 | -0,029297195 | 0 | 0 |
| elbow26 | PS23F | 0,089587504 | 0,089763536 | -0,02304551 | -0,098094267 | 0 | 0 |
| elbow26 | PS3 | -0,005979484 | -0,00598108 | -0,019915376 | -0,029223116 | 0 | 0 |
| elbow26 | PS4 | 0,009056333 | 0,008927168 | -0,017890095 | -0,033233935 | 0 | 0 |
| elbow26 | PS5 | 0,001675416 | 0,001807383 | -0,027746532 | -0,045231685 | 0 | 0 |
| elbow26 | PS6A | -0,016456452 | -0,01648906 | -0,026940778 | -0,033986834 | 0 | 0 |
| elbow26 | PS6B | 0,007897827 | 0,008095195 | -0,015579241 | -0,029229174 | 0 | 0 |
| elbow26 | PS7F | -0,017555353 | -0,017490931 | -0,028004126 | -0,034277634 | 0 | 0 |
| elbow26 | PS9V | -0,002170737 | -0,002279855 | -0,025950991 | -0,041510098 | 0 | 0 |
| elbow26 | PS1 | 0,001568579 | 0,001623625 | -0,013895239 | -0,02387846 | 0 | 0 |
| elbow27 | PS14 | -0,001502453 | -0,001523818 | -0,004122178 | -0,005703728 | 0 | 0 |
| elbow27 | PS15B | -0,002235053 | -0,002225912 | -0,00365519 | -0,004627072 | 0 | 0 |
| elbow27 | PS18C | -0,000739327 | -0,000731078 | -0,004380691 | -0,006782538 | 0 | 0 |
| elbow27 | PS19A | -0,000826989 | -0,000832811 | -0,003397857 | -0,004980939 | 0 | 0 |
| elbow27 | PS19F | 0,001931613 | 0,001993013 | -0,005531714 | -0,009878158 | 0 | 0 |
| elbow27 | PS23F | -0,002876079 | -0,002857505 | -0,004457334 | -0,005481332 | 0 | 0 |
| elbow27 | PS4 | 0,000786925 | 0,000820425 | -0,004477408 | -0,007629195 | 0 | 0 |
| elbow27 | PS5 | -0,002876079 | -0,002897787 | -0,004439 | -0,005296726 | 0 | 0 |
| elbow27 | PS6A | 0,005376712 | 0,005325048 | -0,005275901 | -0,011972059 | 0 | 0 |
| elbow27 | PS6B | -0,002876079 | -0,002876799 | -0,004446053 | -0,005410001 | 0 | 0 |
| elbow27 | PS7F | -0,001367783 | -0,001376371 | -0,004102353 | -0,005898829 | 0 | 0 |
| elbow27 | PS9V | -0,001337618 | -0,001309567 | -0,004214405 | -0,005967961 | 0 | 0 |
| elbow27 | PS1 | 2,63426E-05 | 2,29367E-05 | -0,002415602 | -0,00399632 | 0 | 0 |
| elbow28 | PS14 | -0,005829096 | -0,005841575 | -0,008751248 | -0,01066672 | 0 | 0 |
| elbow28 | PS15B | -0,005829096 | -0,005829722 | -0,008776441 | -0,010623298 | 0 | 0 |
| elbow28 | PS18C | -0,004627173 | -0,004641966 | -0,008176117 | -0,010691942 | 0 | 0 |
| elbow28 | PS19A | 0,001150961 | 0,001181902 | -0,008509404 | -0,014443404 | 0 | 0 |
| elbow28 | PS19F | -0,001512132 | -0,001530297 | -0,00667525 | -0,00997161 | 0 | 0 |
| elbow28 | PS23F | -0,005829096 | -0,005795647 | -0,008838874 | -0,010800972 | 0 | 0 |
| elbow28 | PS3 | 0,013252048 | 0,013414909 | -0,00413123 | -0,014110131 | 0 | 0 |
| elbow28 | PS4 | -0,005829096 | -0,00581431 | -0,008807298 | -0,010717275 | 0 | 0 |
| elbow28 | PS5 | -0,00033459 | -0,000256716 | -0,008848657 | -0,014085374 | 0 | 0 |
| elbow28 | PS6A | -0,004902312 | -0,004929268 | -0,00803739 | -0,010161346 | 0 | 0 |
| elbow28 | PS7F | -0,005829096 | -0,005819174 | -0,008837363 | -0,010691194 | 0 | 0 |
| elbow28 | PS9V | -0,005267614 | -0,005282679 | -0,008466625 | -0,01052424 | 0 | 0 |
| elbow28 | PS1 | 0,003790478 | 0,003757805 | -0,00929694 | -0,017609339 | 0 | 0 |
| elbow29 | PS14 | -0,002140322 | -0,00215041 | -0,007813572 | -0,01123066 | 0 | 0 |
| elbow29 | PS15B | -0,005217246 | -0,005209361 | -0,008334573 | -0,010244502 | 0 | 0 |
| elbow29 | PS18C | -0,001081596 | -0,001068966 | -0,006677747 | -0,010577125 | 0 | 0 |
| elbow29 | PS19A | -0,005217246 | -0,005199245 | -0,008269259 | -0,010159086 | 0 | 0 |
| elbow29 | PS19F | -0,005217246 | -0,005222195 | -0,008358621 | -0,010183833 | 0 | 0 |
| elbow29 | PS23F | 0,005771765 | 0,005844742 | -0,009853394 | -0,019979354 | 0 | 0 |
| elbow29 | PS3 | 0,003365838 | 0,003363335 | -0,005866486 | -0,011290019 | 0 | 0 |
| elbow29 | PS4 | 0,005771765 | 0,005779433 | -0,010484348 | -0,020313164 | 0 | 0 |
| elbow29 | PS5 | 0,002475062 | 0,002498551 | -0,009241623 | -0,016393405 | 0 | 0 |
| elbow29 | PS6A | -0,001554242 | -0,001487983 | -0,00773962 | -0,011793001 | 0 | 0 |
| elbow29 | PS6B | 0,012534234 | 0,012459982 | -0,013051803 | -0,029388595 | 0 | 0 |
| elbow29 | PS7F | -0,005217246 | -0,005217543 | -0,008337411 | -0,010353744 | 0 | 0 |
| elbow29 | PS9V | -0,005217246 | -0,005198385 | -0,008376136 | -0,01025649 | 0 | 0 |
| elbow29 | PS1 | 0,000943723 | 0,001017409 | -0,00501616 | -0,008426932 | 0 | 0 |
| elbow30 | PS14 | -0,007158681 | -0,007215755 | -0,016187593 | -0,021938447 | 0 | 0 |
| elbow30 | PS15B | -0,007158681 | -0,007214552 | -0,016220406 | -0,021588648 | 0 | 0 |
| elbow30 | PS18C | -0,001358925 | -0,001453977 | -0,011798569 | -0,018298165 | 0 | 0 |
| elbow30 | PS19A | -0,000748424 | -0,000735241 | -0,013419275 | -0,021142848 | 0 | 0 |
| elbow30 | PS19F | -0,007158681 | -0,007108113 | -0,016339974 | -0,021587322 | 0 | 0 |
| elbow30 | PS23F | -0,007158681 | -0,007117271 | -0,016243456 | -0,021702009 | 0 | 0 |
| elbow30 | PS3 | -0,006439773 | -0,006427827 | -0,015515565 | -0,021092336 | 0 | 0 |
| elbow30 | PS4 | -0,007158681 | -0,007174904 | -0,016277312 | -0,021854384 | 0 | 0 |

| | | | | | | | |
|---|---|---|---|---|---|---|---|
| elbow30 | PS5 | -0,007158681 | -0,007367158 | -0,016135111 | -0,021278066 | 0 | 0 |
| elbow30 | PS6A | 0,069764396 | 0,071248767 | -0,042847861 | -0,118875207 | 0 | 0 |
| elbow30 | PS6B | 0,002456704 | 0,002553438 | -0,014200821 | -0,02421697 | 0 | 0 |
| elbow30 | PS7F | -0,006404533 | -0,006437813 | -0,015478995 | -0,021321937 | 0 | 0 |
| elbow30 | PS9V | -0,007158681 | -0,007089532 | -0,016387496 | -0,022551909 | 0 | 0 |
| elbow30 | PS1 | -0,007158681 | -0,007124359 | -0,016111016 | -0,022046668 | 0 | 0 |
| elbow31 | PS14 | -0,010730079 | -0,010668044 | -0,019620407 | -0,025330726 | 0 | 0 |
| elbow31 | PS15B | -0,014778662 | -0,014810105 | -0,021113671 | -0,025266382 | 0 | 0 |
| elbow31 | PS18C | -0,006462763 | -0,00650506 | -0,016517703 | -0,023100924 | 0 | 0 |
| elbow31 | PS19A | -0,014778662 | -0,01477139 | -0,021078242 | -0,025169041 | 0 | 0 |
| elbow31 | PS19F | -0,006876564 | -0,00698061 | -0,017923216 | -0,024623922 | 0 | 0 |
| elbow31 | PS23F | 0,042913645 | 0,043111065 | -0,018471824 | -0,058837117 | 0 | 0 |
| elbow31 | PS4 | -0,007785655 | -0,007962299 | -0,018892927 | -0,025900175 | 0 | 0 |
| elbow31 | PS5 | -0,007785655 | -0,007769632 | -0,019384727 | -0,025835026 | 0 | 0 |
| elbow31 | PS6A | -0,001958149 | -0,001915812 | -0,021230405 | -0,033000574 | 0 | 0 |
| elbow31 | PS6B | 0,011742893 | 0,011697777 | -0,010060187 | -0,024487232 | 0 | 0 |
| elbow31 | PS7F | -0,014778662 | -0,014772149 | -0,021270608 | -0,025128847 | 0 | 0 |
| elbow31 | PS9V | 0,004102457 | 0,004013069 | -0,018828588 | -0,034066119 | 0 | 0 |
| elbow31 | PS1 | 0,008952762 | 0,008988231 | -0,008469589 | -0,019775613 | 0 | 0 |
| elbow32 | PS14 | -0,004009024 | -0,004006343 | -0,006990242 | -0,008852615 | 0 | 0 |
| elbow32 | PS15B | -0,004009024 | -0,004034052 | -0,006855584 | -0,008659844 | 0 | 0 |
| elbow32 | PS18C | 0,02355165 | 0,023456418 | -0,003337296 | -0,018773504 | 0 | 0 |
| elbow32 | PS19A | -0,004009024 | -0,004014748 | -0,006960654 | -0,008767757 | 0 | 0 |
| elbow32 | PS19F | -0,004009024 | -0,004012184 | -0,007019053 | -0,008830223 | 0 | 0 |
| elbow32 | PS23F | -0,004009024 | -0,004017442 | -0,006938623 | -0,008692345 | 0 | 0 |
| elbow32 | PS3 | -0,002473271 | -0,002444306 | -0,005868497 | -0,007870452 | 0 | 0 |
| elbow32 | PS4 | -0,004009024 | -0,004017451 | -0,00698024 | -0,008885097 | 0 | 0 |
| elbow32 | PS5 | 0,003683283 | 0,003752899 | -0,007664058 | -0,014624295 | 0 | 0 |
| elbow32 | PS6A | -0,004009024 | -0,004005558 | -0,006950535 | -0,008729498 | 0 | 0 |
| elbow32 | PS6B | -0,004009024 | -0,004017925 | -0,006960639 | -0,008869954 | 0 | 0 |
| elbow32 | PS7F | -0,004009024 | -0,003995287 | -0,006969606 | -0,008788513 | 0 | 0 |
| elbow32 | PS9V | -0,004009024 | -0,004003561 | -0,006958245 | -0,008664635 | 0 | 0 |
| elbow32 | PS1 | 0,015328582 | 0,015319149 | -0,000558486 | -0,011090123 | 0 | 0 |
| elbow33 | PS14 | -0,004458527 | -0,004446506 | -0,00764191 | -0,009696976 | 0 | 0 |
| elbow33 | PS15B | -0,004458527 | -0,004460602 | -0,00764936 | -0,00962458 | 0 | 0 |
| elbow33 | PS18C | 0,009116133 | 0,009149549 | -0,010759172 | -0,02365769 | 0 | 0 |
| elbow33 | PS19A | -0,004458527 | -0,004452224 | -0,007634145 | -0,009593916 | 0 | 0 |
| elbow33 | PS19F | -0,001381604 | -0,001346135 | -0,006433156 | -0,009507685 | 0 | 0 |
| elbow33 | PS23F | -0,004458527 | -0,004474801 | -0,007642066 | -0,009546894 | 0 | 0 |
| elbow33 | PS3 | 0,003218731 | 0,003168378 | -0,005349912 | -0,010478812 | 0 | 0 |
| elbow33 | PS4 | -0,004458527 | -0,004464052 | -0,00757389 | -0,009663072 | 0 | 0 |
| elbow33 | PS5 | -0,002627026 | -0,002675878 | -0,006350963 | -0,008705816 | 0 | 0 |
| elbow33 | PS6A | -0,004458527 | -0,00450159 | -0,007539713 | -0,009421045 | 0 | 0 |
| elbow33 | PS7F | -0,004458527 | -0,004447223 | -0,007549511 | -0,009518906 | 0 | 0 |
| elbow33 | PS9V | -0,004458527 | -0,004456104 | -0,007663481 | -0,009562799 | 0 | 0 |
| elbow33 | PS1 | 0,001951729 | 0,001916146 | -0,004809794 | -0,009100864 | 0 | 0 |
| elbow34 | PS14 | -0,006176446 | -0,006204288 | -0,012639172 | -0,017008793 | 0 | 0 |
| elbow34 | PS15B | -0,009253369 | -0,009231446 | -0,013684282 | -0,016396267 | 0 | 0 |
| elbow34 | PS18C | 0,001880448 | 0,001900289 | -0,006162249 | -0,01157847 | 0 | 0 |
| elbow34 | PS19A | -0,009253369 | -0,009273382 | -0,013661125 | -0,016507389 | 0 | 0 |
| elbow34 | PS19F | 0,016106356 | 0,016015546 | -0,005678195 | -0,019226875 | 0 | 0 |
| elbow34 | PS23F | -0,007116617 | -0,007152037 | -0,012861974 | -0,016384336 | 0 | 0 |
| elbow34 | PS3 | -0,008837569 | -0,008795815 | -0,013326855 | -0,016274729 | 0 | 0 |
| elbow34 | PS4 | -0,002260362 | -0,002169504 | -0,013753094 | -0,020913496 | 0 | 0 |
| elbow34 | PS5 | 0,008728649 | 0,009037606 | -0,010934784 | -0,022627755 | 0 | 0 |
| elbow34 | PS6A | -0,009253369 | -0,00930375 | -0,013654433 | -0,016366949 | 0 | 0 |
| elbow34 | PS6B | 0,042761283 | 0,042654552 | -0,000318245 | -0,027679148 | 0 | 0 |
| elbow34 | PS7F | -0,009253369 | -0,00925582 | -0,013683489 | -0,016644716 | 0 | 0 |
| elbow34 | PS9V | -0,009253369 | -0,009223693 | -0,013841916 | -0,016461245 | 0 | 0 |
| elbow34 | PS1 | 0,001181104 | 0,001170596 | -0,006907386 | -0,011511939 | 0 | 0 |
| elbow35 | PS14 | -0,015730517 | -0,015664799 | -0,024808738 | -0,030422804 | 0 | 0 |
| elbow35 | PS15B | -0,012086792 | -0,012212054 | -0,026441219 | -0,03606272 | 0 | 0 |
| elbow35 | PS19A | 0,005861926 | 0,006098181 | -0,022515663 | -0,039951599 | 0 | 0 |
| elbow35 | PS19F | -0,012786093 | -0,012831423 | -0,024358727 | -0,03203218 | 0 | 0 |
| elbow35 | PS23F | 0,018682439 | 0,019001778 | -0,019205151 | -0,04231492 | 0 | 0 |
| elbow35 | PS3 | 0,011119407 | 0,01125888 | -0,009715274 | -0,022951481 | 0 | 0 |
| elbow35 | PS4 | -0,0197791 | -0,019663918 | -0,027550622 | -0,032055763 | 0 | 0 |
| elbow35 | PS5 | -0,0197791 | -0,019745503 | -0,027393235 | -0,032096029 | 0 | 0 |
| elbow35 | PS6A | -0,014971407 | -0,014950319 | -0,024526881 | -0,030378158 | 0 | 0 |
| elbow35 | PS7F | -0,0197791 | -0,019775448 | -0,027570229 | -0,032223643 | 0 | 0 |

| | | | | | | | |
|---|---|---|---|---|---|---|---|
| elbow35 | PS9V | 0,002392846 | 0,002356738 | -0,021315838 | -0,036024219 | 0 | 0 |
| elbow35 | PS1 | 0,013678878 | 0,013735151 | -0,012583486 | -0,029294312 | 0 | 0 |

**Supplementary table 2. Statistical results from linear regression model bootstrapping for Spn cross-sectional phenotype and serotype analysis.** The observed (obs_estimate) and estimated (unbiased_estimate) proportions of clusters among total B cells for a given serotype-specificity and sample, were generated with an upper-sided linear regression model with wild bootstrap simulation (9999 resamples). Confidence intervals (ci) and Bonferroni's method adjusted ci (ci.adj) are shown, as well as statistical significance (signif_ci/ signif_ci.adj = 1).

| Supplementary table 3 | | | | | | | |
|---|---|---|---|---|---|---|---|
| cluster | timepoint | obs_estimate | unbiased_estimate | ci_limit_0.05 | ci.adj_limit_0.025 | signif_ci | signif_ci.adj |
| elbow07 | M4 | 0.0387190677018709 | 0.0388413938599702 | 0.018539172120877 | 0.0146486905440044 | 1 | 1 |
| elbow11 | M4 | 0.0454526765194477 | 0.0454197192337327 | 0.0187444485062122 | 0.0132632268303365 | 1 | 1 |
| elbow03 | M4 | 0.095696755686353 | 0.0960637957209242 | 0.0520957451537589 | 0.0445783096409878 | 1 | 1 |
| elbow09 | M4 | 0.0125221006471866 | 0.0124811661635298 | 0.00513192131803552 | 0.00389584391165417 | 1 | 1 |
| elbow13 | M4 | 0.00615223040135064 | 0.00613942635972125 | 0.00253597562536561 | 0.00184346800988204 | 1 | 1 |
| elbow22 | M4 | 0.0201158966958282 | 0.0201486626538739 | 0.00809270135571964 | 0.00603222312614188 | 1 | 1 |
| elbow05 | M4 | 0.00907469424297034 | 0.00906400285652889 | -0.000373093228988612 | -0.00225446084750009 | 0 | 0 |
| elbow06 | M4 | 0.0180891120160918 | 0.0181043629341241 | -0.000036215036735241 | -0.0034584209879452 | 0 | 0 |
| elbow01 | M4 | -0.0024981382599618 | -0.0024132966033086 | -0.0132151651199272 | -0.0152337353058856 | 0 | 0 |
| elbow02 | M4 | 0.0130494227051925 | 0.0131843989380414 | -0.0097832443004755 | -0.0142626183174471 | 0 | 0 |
| elbow04 | M4 | 0.00933073584338708 | 0.00938929388130626 | -0.0416823581153119 | -0.0513514973457047 | 0 | 0 |
| elbow08 | M4 | -0.00513281401185176 | -0.0051082719673856 | -0.019118891888884 | -0.021894343648739 | 0 | 0 |
| elbow10 | M4 | -0.0231473980815505 | -0.022952299350188 | -0.0431189156675798 | -0.0469479950871325 | 0 | 0 |
| elbow12 | M4 | 0.00644912123054663 | 0.00654913946661539 | -0.00255756237309722 | -0.00412868342618228 | 0 | 0 |
| elbow14 | M4 | -0.00761641811329294 | -0.00765142033868941 | -0.0160808366775275 | -0.0178036956686031 | 0 | 0 |
| elbow15 | M4 | 0.00355389260749793 | 0.00358463673558799 | -0.000961425652625518 | -0.00186405755249227 | 0 | 0 |
| elbow16 | M4 | 0.0132181150025836 | 0.013311303824617 | -0.00207482636801922 | -0.00495417305237344 | 0 | 0 |
| elbow17 | M4 | -0.00474035768153415 | -0.00469576377506566 | -0.011455676010764 | -0.012707994428801 | 0 | 0 |
| elbow18 | M4 | 0.000183150183150183 | 0.00020787500969714 | -0.00686073902871814 | -0.00811112572691376 | 0 | 0 |
| elbow19 | M4 | -0.00264732205181388 | -0.00265344592169131 | -0.00760313084755765 | -0.00856506241830524 | 0 | 0 |
| elbow20 | M4 | -0.00624627664103402 | -0.00633196374608449 | -0.0156518562839745 | -0.0173514161278738 | 0 | 0 |
| elbow21 | M4 | -0.020435225185977 | -0.0205753313048421 | -0.038636419949816 | -0.0418786441611714 | 0 | 0 |
| elbow23 | M4 | -0.00160144953991246 | -0.00157102123058828 | -0.00598616834210475 | -0.00681272210889326 | 0 | 0 |
| elbow24 | M4 | -0.00653317402456673 | -0.00653876520719599 | -0.0147338873531064 | -0.0163171306459282 | 0 | 0 |
| elbow25 | M4 | -0.00811143818790515 | -0.00802723956142462 | -0.0232008980978041 | -0.025933154315713 | 0 | 0 |
| elbow26 | M4 | -0.0207794000416989 | -0.0206931481316721 | -0.0415914211065924 | -0.0456373908397772 | 0 | 0 |
| elbow27 | M4 | 0.000616279268627808 | 0.000638946994774594 | -0.00219779376716285 | -0.0027438291857353 | 0 | 0 |
| elbow28 | M4 | -0.00439108075221043 | -0.00442172252036754 | -0.0102328711588892 | -0.0112743278210812 | 0 | 0 |
| elbow29 | M4 | -0.003956830616703 | -0.0039928514185696 | -0.00977113592871606 | -0.0108868166055653 | 0 | 0 |
| elbow30 | M4 | -0.0126474625316218 | -0.0128184683128676 | -0.0301570108575694 | -0.0335984959939659 | 0 | 0 |
| elbow31 | M4 | 0.00681544093296104 | 0.00683167417119726 | -0.0108668491794457 | -0.0141446526499419 | 0 | 0 |
| elbow32 | M4 | -0.00771034257342575 | -0.00771362103385678 | -0.0130500235349328 | -0.0140847556191554 | 0 | 0 |
| elbow33 | M4 | -0.00891705485092482 | -0.00890945486128366 | -0.0144818411428929 | -0.0154814990882119 | 0 | 0 |
| elbow34 | M4 | -0.00612131899947026 | -0.00615487141352728 | -0.0159743002418259 | -0.017846783846472 | 0 | 0 |
| elbow35 | M4 | -0.0221788159132166 | -0.0221201276911835 | -0.0353166393577675 | -0.037693740775137 | 0 | 0 |

The observed (obs_estimate) and estimated (unbiased_estimate) difference in proportion of clusters among total B cells comparing PCV13-vaccinated (M4) to non-vaccinated individuals were generated with an upper-sided linear regression model with wild bootstrap simulation (9999 resamples). Confidence intervals (ci) and Bonferroni's method adjusted ci (ci.adj) are shown, as well as statistical significance (signif_ci/ signif_ci.adj = 1).

| Supplementary table 4 | | | | | | | |
|---|---|---|---|---|---|---|---|
| **Marker/dye** | **Fluorochrome** | **Clone** | **Isotype** | **Company** | **Catalogue** | **Dilution** | **Surface/intracel.** |
| CD7 | BV510 | M-T701 | Mouse IgG1. κ | BD | 563650 | 100 | Surface |
| CD19 | BUV395 | HIB19 | Mouse IgG1. κ | BD | 740287 | 200 | Surface |
| HLA-DR | BUV496 | G46-6 | Mouse IgG2a. κ | BD | 749866 | 200 | Surface |
| IgD | BUV563 | I-A6-2 | Mouse IgG2a. κ | BD | 741394 | 400 | Surface |
| IgM | BV570 | MHM-88 | Mouse IgG1. κ | Biolegend | 314517 | 400 | Surface |
| IgA | APC-Vio770 | IS11-8E10 | Mouse IgG1κ | Miltenyi | 130-113-999 | 1600 | Surface |
| IgG | PE-CF594 | G18-145 | Mouse IgG1. κ | BD | 562538 | 400 | Surface |
| CD38 | APC-Fire810 | HIT2 | Mouse IgG1. κ | Biolegend | 303550 | 100 | Surface |
| CD10 | BV605 | HI 10 a | Mouse IgG1. κ | Biolegend | 312222 | 100 | Surface |
| CD11c | BV650 | Bu15 | Mouse IgG1. κ | Biolegend | 337237 | 400 | Surface |
| CD20 | Pacific orange | HI47 | Mouse IgG3 | ThermoFisher | MHCD2030 | 20 | Surface |
| CD21 | BUV805 | BLy4 | Mouse IgG1. κ | BD | 742008 | 200 | Surface |
| CD27 | APC-R700 | M-T271 | Mouse IgG1. κ | BD | 565116 | 100 | Surface |
| CD43 | PerCP-Cy5.5 | 1G10 | Mouse IgG1. κ | BD | 563521 | 400 | Surface |
| $CD45RB_{MEM55}$ | PE | MEM-55 | Mouse IgG2b. κ | Biolegend | 310204 | 400 | Surface |
| CD5 | BV750 | L17F12 | Mouse IgG2a. κ | BD | 747090 | 200 | Surface |
| CD73 | AF647 | AD2 | Mouse IgG1 | Abcam | 243083 | 200 | Surface |
| CD95 | PE-Cy5 | DX2 | Mouse IgG1. κ | Biolegend | 305610 | 400 | Surface |
| CD80 | PerCP-eFluor710 | 16-10A1 | Hamster IgG | ThermoFisher | 46-0801-82 | 200 | Surface |
| CXCR3 | PE-Cy7 | G025H7 | Mouse IgG1. κ | Biolegend | 353719 | 1200 | Surface |
| PD-1 | BV480 | EH12.1 | Mouse IgG1. κ | BD | 566112 | 100 | Surface |
| LAIR1 | BUV737 | DX26 (RUO) | Mouse IgG1. κ | BD | 749446 | 400 | Surface |
| IRF4 | APC | REA201 | Human IgG1 | Miltenyi | 130-100-915 | 200 | Intracellular |
| Ki-67 | BV711 | Ki67 | Mouse IgG1. κ | Biolegend | 350515 | 400 | Intracellular |
| T-bet | BV785 | 4B10 | Mouse IgG1. κ | Biolegend | 644835 | 100 | Intracellular |
| SA-BB515 | BB515 | NA | NA | BD | 564453 | NA | NA |
| SA-BUV615 | BUV615 | NA | NA | BD | 613013 | NA | NA |
| SA-BUV661 | BUV661 | NA | NA | BD | 612979 | NA | NA |
| SA-BV421 | BV421 | NA | NA | BD | 563259 | NA | NA |
| Live/Dead | Blue | NA | NA | ThermoFisher | L34962 | 500 | NA |

**Supplementary table 4. Antibody list for GBS analysis.**

| Supplementary table 5 | | | | | | | | | |
|---|---|---|---|---|---|---|---|---|---|
| **cluster** | **type** | **obs_estimate** | **unbiased_estimate** | **ci_limit_0.025** | **ci_limit_0.975** | **ci.adj_limit_0.005** | **ci.adj_limit_0.995** | **signif_ci** | **signif_ci.adj** |
| IgG | PSIa | 0.297912206130363 | 0.299039856317133 | 0.14975417051434 | 0.439840962827398 | 0.105861315750359 | 0.484584723888382 | 1 | 1 |
| IgG | PSIII | 0.120990436331345 | 0.120348081226637 | 0.00512145527064722 | 0.236209174402328 | -0.0307200461421479 | 0.272579090230043 | 1 | 0 |
| IgA | PSIb | 0.00374230270620651 | 0.0041038714582251 | -0.0731874274227158 | 0.0784250252830294 | -0.0981078742549699 | 0.10619671130899 | 0 | 0 |
| IgA | PSII | 0.0282544701118572 | 0.0280538287595615 | -0.105233745733272 | 0.16115871866462 | -0.145758466096581 | 0.20240877434359 | 0 | 0 |
| IgA | PSIII | 0.0174878782833328 | 0.0175517034327768 | -0.0504831232030867 | 0.0837804102267365 | -0.0723371707456517 | 0.105155227086669 | 0 | 0 |
| IgA | PSV | 0.0590732531357532 | 0.0589324807687921 | -0.00488407139076333 | 0.12280419906185 | -0.0271181827890558 | 0.142750081950293 | 0 | 0 |
| IgA | PSIa | 0.0825265914546965 | 0.0825261948947213 | -0.0208315201777741 | 0.186392211228752 | -0.0533861301148082 | 0.219659165429992 | 0 | 0 |
| IgD | PSIb | 0.114279492788851 | 0.114999559615973 | -0.0254726331380426 | 0.252777487961871 | -0.0714173612445071 | 0.298212952356342 | 0 | 0 |
| IgD | PSII | 0.0804357654532321 | 0.0803485784807978 | -0.0343924637993539 | 0.192919831970397 | -0.0668466782092957 | 0.228838275655617 | 0 | 0 |
| IgD | PSIII | -0.0010630152675607 | -0.00148774520505988 | -0.242029634851683 | 0.241140763767503 | -0.318355949156198 | 0.319648596447281 | 0 | 0 |
| IgD | PSV | -0.086894529082029 | -0.0863578701529576 | -0.236582454662417 | 0.0608480849133081 | -0.28465198825595 | 0.111418006077663 | 0 | 0 |
| IgD | PSIa | -0.210516905278866 | -0.208617418338795 | -0.457612174854376 | 0.027181080633052 | -0.544242098592279 | 0.104652171987272 | 0 | 0 |
| IgDIgM | PSIb | 0.230801328795981 | 0.231161286988667 | -0.00438615519772776 | 0.462282258793965 | -0.0765229108043725 | 0.544048137690967 | 0 | 0 |
| IgDIgM | PSII | 0.0603747901295082 | 0.0599010533399151 | -0.0817671783205953 | 0.202582014900993 | -0.126428528093155 | 0.246201703592698 | 0 | 0 |
| IgDIgM | PSIII | 0.0917518087972635 | 0.0949304976525217 | -0.143318905752151 | 0.31734454092602 | -0.209255963778439 | 0.389874649361361 | 0 | 0 |
| IgDIgM | PSV | -0.0153449328449335 | -0.0156603894889066 | -0.227378714682933 | 0.19683138191507 | -0.297455550715123 | 0.260827421255727 | 0 | 0 |
| IgDIgM | PSIa | -0.0480003198516886 | -0.0471670132007757 | -0.234020395465675 | 0.136497218184659 | -0.295560259870355 | 0.189966340780082 | 0 | 0 |
| IgG | PSIb | 0.0379203127532006 | 0.0376857804751357 | -0.137550216409217 | 0.212923221276193 | -0.195321775849025 | 0.266616318800949 | 0 | 0 |
| IgG | PSII | -0.135219839626075 | -0.135009395399472 | -0.354144100924798 | 0.0809770097187282 | -0.429366943269422 | 0.148928214196056 | 0 | 0 |
| IgG | PSV | 0.0683729464979467 | 0.067544559033488 | -0.0664245684036486 | 0.202600617264328 | -0.103455824488959 | 0.241166797812673 | 0 | 0 |
| IgM | PSIb | -0.10265252795333 | -0.102201983685883 | -0.2969608707855 | 0.0898658093530763 | -0.35449524937631 | 0.153306546594738 | 0 | 0 |
| IgM | PSII | -0.0338451860685226 | -0.0337354451266717 | -0.136917589525561 | 0.067127647711779 | -0.167314891572706 | 0.0979144678319177 | 0 | 0 |
| IgM | PSIII | -0.0246216535989263 | -0.0247651979896518 | -0.13063977350382 | 0.0812224856260448 | -0.162315552306208 | 0.114364334802359 | 0 | 0 |
| IgM | PSV | -0.0252067377067377 | -0.0255429139515515 | -0.190976250204212 | 0.135259826247834 | -0.242361959797995 | 0.184721834662163 | 0 | 0 |
| IgM | PSIa | -0.121921572454504 | -0.120939621350069 | -0.310653979024049 | 0.0655352417517152 | -0.379912991307615 | 0.126446504778936 | 0 | 0 |

**Supplementary table 5. Statistical results from linear regression model bootstrapping for GBS isotype and population analysis.** Table shows the observed (obs_estimate) and estimated (unbiased_estimate) difference in proportions of clusters among total B cells for a given serotype-specificity and B cell isotype, comparing South African- with Dutch donors. The analysis was performed using a two-sided linear regression model with wild bootstrap simulation (9999 resamples). Confidence intervals (ci) and Bonferroni's method adjusted ci (ci.adj) are shown, as well as statistical significance (signif_ci/ signif_ci.adj = 1).

| Phenotype and population analysis | | | | | | | | | |
|---|---|---|---|---|---|---|---|---|---|
| cluster | type | obs_estimate | unbiased_estimate | ci_limit_0.025 | ci_limit_0.975 | ci.adj_limit_0.005 | ci.adj_limit_0.995 | signif_ci | signif_ci.adj |
| elbow06 | PSIb | 1,08E-03 | 1,08E-03 | 6,56E-04 | 1,50E-03 | 5,14E-04 | 1,63E-04 | 1 | 1 |
| elbow14 | PSII | 0.0126875708961271 | 0.0127517387360139 | 0.00342290586185557 | 0.0216497293647453 | 0.0004951804991052 | 0.0249148379803903 | 1 | 1 |
| elbow20 | PSIa | 0.039257444361733 | 0.0391506993020294 | 0.0106205177344011 | 0.0681754258113157 | 0.00146429143611112 | 0.0779807106691606 | 1 | 1 |
| elbow31 | PSIa | 0.0601316268135356 | 0.0601352378480176 | 0.0220846436838107 | 0.0985433461820823 | 0.00878887268485867 | 0.11032963397321 | 1 | 1 |
| elbow31 | PSV | 0.0666205757332066 | 0.0665026687064256 | 0.03271904742664 | 0.100336792367083 | 0.0225189647807544 | 0.111336940022469 | 1 | 1 |
| elbow22 | PSIa | 0.066972619152292 | 0.0669317225693528 | 0.025005414689214 | 0.109269225564805 | 0.0130963451706423 | 0.122093372752177 | 1 | 1 |
| elbow26 | PSV | 0.0774340381723222 | 0.077346458162919 | 0.0272364866545509 | 0.127828463131688 | 0.0104859658561076 | 0.142727641269059 | 1 | 1 |
| elbow24 | PSIa | 0.126869289581038 | 0.127306786528142 | 0.0692927514343191 | 0.183538832522994 | 0.0508724214131485 | 0.201947525137912 | 1 | 1 |
| elbow09 | PSV | -1,80E-02 | -1,80E-02 | -3,05E-02 | -5,55E-03 | -3,47E-02 | 0 | 1 | 0 |
| elbow05 | PSIII | -3,47E-03 | -3,46E-03 | -1,18E-02 | 5,55E-03 | -1,39E-02 | 8,33E-03 | 0 | 0 |
| elbow12 | PSV | -2,08E-03 | -2,10E-03 | -6,96E-03 | 2,92E-03 | -8,41E-04 | 4,44E-06 | 0 | 0 |
| elbow17 | PSV | -1,04E-03 | -1,02E-04 | -5,20E-03 | 2,78E-03 | -6,25E-03 | 4,16E-03 | 0 | 0 |
| elbow11 | PSV | -3,47E-04 | -3,44E-04 | -1,83E-03 | 1,13E-03 | -2,27E-03 | 1,65E-03 | 0 | 0 |
| elbow01 | PSIb | -6,41E-07 | -6,40E-05 | -5,83E-04 | 4,49E-05 | -7,43E-04 | 5,82E-04 | 0 | 0 |
| elbow28 | PSIb | 0 | -1,72E-06 | -2,78E-03 | 2,78E-03 | -2,78E-03 | 2,78E-03 | 0 | 0 |
| elbow01 | PSII | 4,47E-04 | 4,56E-04 | -2,04E-03 | 2,93E-03 | -2,78E-04 | 3,85E-04 | 0 | 0 |
| elbow11 | PSIb | 6,07E-04 | 6,10E-04 | -1,81E-04 | 1,41E-03 | -4,63E-04 | 1,79E-03 | 0 | 0 |
| elbow17 | PSIb | 6,94E-04 | 7,04E-04 | -2,08E-03 | 3,47E-03 | -2,78E-03 | 4,17E-03 | 0 | 0 |
| elbow17 | PSIII | 1,39E-03 | 1,43E-03 | -2,78E-03 | 5,55E-03 | -4,16E-03 | 6,94E-03 | 0 | 0 |
| elbow01 | PSIa | 1,69E-03 | 1,70E-03 | -2,15E-03 | 5,43E-03 | -3,13E-03 | 6,60E-03 | 0 | 0 |
| elbow06 | PSII | 0.000478468899521532 | 0.000485410092788471 | -0.000447147051866367 | 0.0013748245556312 | -0.000712135168113252 | 0.00164772267476296 | 0 | 0 |
| elbow22 | PSII | -0.00052005012531324 | -0.00068056501808971 | -0.0241004618168165 | 0.0240359823905704 | -0.0311660547254018 | 0.0326140042048083 | 0 | 0 |
| elbow15 | PSII | -0.00085668717247663 | -0.00076680800103428 | -0.0110940669193567 | 0.00917861826597592 | -0.014281100689971 | 0.0121335324869471 | 0 | 0 |
| elbow05 | PSII | 0.000909090909090839 | 0.000924093765248404 | -0.000904768802075086 | 0.00258900264894773 | -0.00141364336287523 | 0.00314597773882602 | 0 | 0 |
| elbow30 | PSIa | -0.00104249284645657 | -0.0012778953184661 | -0.0659383353503506 | 0.0656609612462084 | -0.0855536624066189 | 0.0885150601827155 | 0 | 0 |
| elbow10 | PSII | 0.00110951687569244 | 0.0011871449268079 | -0.0978012740250651 | 0.101531340852114 | -0.128677069884458 | 0.131719610642969 | 0 | 0 |
| elbow18 | PSII | 0.00113458097409205 | 0.000429603448215514 | -0.07812309233905 | 0.0830111531312705 | -0.101965575159898 | 0.106685172430412 | 0 | 0 |
| elbow07 | PSIa | 0.00126262626262622 | 0.00125447999277994 | -0.00115595771128142 | 0.00368641200139634 | -0.00189145098812117 | 0.00446348561002362 | 0 | 0 |
| elbow03 | PSV | 0.00154401154401175 | 0.0015964784304912 | -0.0230261447620116 | 0.0252524353547185 | -0.030111609579553 | 0.0328407172140487 | 0 | 0 |
| elbow06 | PSIa | 0.00184275184275202 | 0.00184070311152699 | -0.00168249573099555 | 0.00538649396411933 | -0.00264907931558245 | 0.00641960730848161 | 0 | 0 |
| elbow30 | PSII | -0.0018553828665002 | -0.00185043208295685 | -0.0488014166649993 | 0.0448501024610332 | -0.0621865661815629 | 0.0585805237596373 | 0 | 0 |
| elbow07 | PSV | 0.00192722681359046 | 0.00198820533949921 | -0.0239868482271311 | 0.0278874352556704 | -0.0321503777791115 | 0.0363675108838208 | 0 | 0 |
| elbow15 | PSIb | 0.00220959595959597 | 0.00222336804002774 | -0.0101978238106322 | 0.0149557898081323 | -0.0143337479123241 | 0.0186850980529172 | 0 | 0 |
| elbow29 | PSII | -0.00221104324511935 | -0.0023453768767322 | -0.0623444828076246 | 0.0589054594023438 | -0.0803848486320499 | 0.0775399244787909 | 0 | 0 |
| elbow11 | PSIII | 0.00267379679144385 | 0.00267374062219731 | -0.00236695793423428 | 0.00788816200348681 | -0.00389574857612204 | 0.00939581257280887 | 0 | 0 |
| elbow14 | PSV | 0.00267379679144386 | 0.00267706789357653 | -0.00244734860451049 | 0.00773878486007679 | -0.00383318614274838 | 0.00921245090035603 | 0 | 0 |
| elbow31 | PSII | 0.00269981075390453 | 0.00236408935424468 | -0.101302299478061 | 0.105782497642674 | -0.132265709404823 | 0.142099665410648 | 0 | 0 |
| elbow14 | PSIa | 0.00284090909090907 | 0.00282773945858111 | -0.00256894876971969 | 0.00840654004993065 | -0.00419129550625583 | 0.00997256065589062 | 0 | 0 |
| elbow22 | PSIII | 0.00287548903859058 | 0.00264433164638064 | -0.0402765768224811 | 0.0470387887386465 | -0.0545135544310753 | 0.0606215738828322 | 0 | 0 |
| elbow09 | PSII | 0.00297619047619034 | 0.00298412123538537 | -0.0134845480028968 | 0.0197025616997223 | -0.0184149737563093 | 0.0247748710084571 | 0 | 0 |
| elbow30 | PSIII | 0.00299586776859505 | 0.00303748366785688 | -0.0457418766935958 | 0.0513857395076625 | -0.0616753938877363 | 0.0659534516752862 | 0 | 0 |
| elbow33 | PSIII | 0.00303030303030307 | 0.00305561151553908 | -0.00275880723366494 | 0.00890176055702721 | -0.00455606778983024 | 0.0106419960368761 | 0 | 0 |
| elbow33 | PSV | 0.00331611570247927 | 0.0033298658817477 | -0.0260497926599029 | 0.033141352252703 | -0.0364450234644205 | 0.0429321567424798 | 0 | 0 |
| elbow05 | PSIb | 0.00349650349650347 | 0.00353255278918301 | -0.00328600022965457 | 0.010035063548881 | -0.00529008907365536 | 0.0123001863767965 | 0 | 0 |
| elbow11 | PSII | -0.00353372434017594 | -0.0035795028619778 | -0.0130348414204964 | 0.00621985493383202 | -0.0156895148979071 | 0.00904174856165367 | 0 | 0 |
| elbow25 | PSII | 0.0038277511961723 | 0.00379890470792276 | -0.00350279029476001 | 0.0110788188421384 | -0.00559045113282623 | 0.0132077484254322 | 0 | 0 |
| elbow23 | PSIa | 0.00397515206887845 | 0.00393662143659898 | 0.000626920095460139 | 0.007417112453184 | -0.000458401528310861 | 0.00861303172213742 | 1 | 0 |
| elbow03 | PSIII | 0.00413223140495864 | 0.00410309650839074 | -0.00375477237741979 | 0.0121494890228649 | -0.00624765777674245 | 0.014589541058429 | 0 | 0 |
| elbow28 | PSII | 0.0044111378152125 | 0.00451022926722728 | -0.0134915009768299 | 0.0220528093375507 | -0.0189982377323745 | 0.0273561510404996 | 0 | 0 |
| elbow19 | PSIa | 0.00452280228674301 | 0.00457668583412046 | -0.00153188171128287 | 0.0105342889673481 | -0.00336522832997142 | 0.0126062700556459 | 0 | 0 |
| elbow19 | PSV | -0.005 | -0.00500912728711257 | -0.0143076974084012 | 0.00406847627438736 | -0.0176612003846256 | 0.00673710026513987 | 0 | 0 |
| elbow32 | PSIa | 0.00501067341517255 | 0.00500963964123243 | -0.0020272102781133 | 0.0119844785927089 | -0.00410860735714855 | 0.0143952696765176 | 0 | 0 |
| elbow16 | PSIb | 0.00505050505050505 | 0.00509418304554701 | -0.0045793057259238 | 0.0147070477324004 | -0.00777098838928628 | 0.0178154629152215 | 0 | 0 |
| elbow23 | PSIb | 0.00505050505050505 | 0.00502224010273269 | -0.00443732443843088 | 0.0146172176167119 | -0.00748407170775788 | 0.0176962632109214 | 0 | 0 |
| elbow18 | PSIb | 0.00505050505050506 | 0.00509001321846675 | -0.00469020905574599 | 0.0147099020576005 | -0.00775966423869812 | 0.0177401275708017 | 0 | 0 |
| elbow17 | PSII | 0.00542699724517908 | 0.0054843514631355 | -0.00114258676448865 | 0.0118388205605084 | -0.0033430131066615 | 0.0140366863934024 | 0 | 0 |
| elbow03 | PSIb | 0.00561868686868676 | 0.00572922945750138 | -0.0130158522555559 | 0.0237187051781912 | -0.0191694053583803 | 0.0296381096555069 | 0 | 0 |
| elbow09 | PSIb | 0.00568181818181808 | 0.00570156977700069 | -0.00515103294613404 | 0.0166886256597083 | -0.00836744709994093 | 0.020204810794621 | 0 | 0 |
| elbow08 | PSV | 0.00568181818181819 | 0.00570870689656877 | -0.00515138402408402 | 0.0162666347480293 | -0.00845147990948512 | 0.0198416139626872 | 0 | 0 |
| elbow25 | PSIa | 0.00575232164799906 | 0.00574159167700049 | -0.00166168728935898 | 0.0131793758746072 | -0.00417025686009526 | 0.0155416427649207 | 0 | 0 |
| elbow20 | PSIII | 0.0058618075329307 | 0.00534787110490819 | -0.157295394618755 | 0.170189447940996 | -0.20413806044395 | 0.22790092050609 | 0 | 0 |
| elbow15 | PSV | 0.00603109012199924 | 0.00598260110243875 | -0.000778006862672829 | 0.0129096913012711 | -0.00284042842754717 | 0.0152210801066907 | 0 | 0 |
| elbow19 | PSII | 0.00623179850452577 | 0.00619747092939639 | -0.0108302659194331 | 0.0232678046971214 | -0.0157037256159318 | 0.0289141146719326 | 0 | 0 |
| elbow12 | PSIII | 0.00649350649350648 | 0.00653826204011345 | -0.0059472048997847 | 0.0188494787389582 | -0.00937589910724547 | 0.0227957897551866 | 0 | 0 |
| elbow07 | PSIII | 0.00649350649350649 | 0.00659296975788493 | -0.00601898186718304 | 0.0188541360783645 | -0.00980851550954934 | 0.0229132494090983 | 0 | 0 |
| elbow11 | PSIa | 0.00660633873091212 | 0.00666023227836503 | -0.00444256345377063 | 0.017341815146621 | -0.00771815944207531 | 0.0210555308488527 | 0 | 0 |
| elbow10 | PSV | -0.00703041393083252 | -0.00712834279471148 | -0.0757013057788884 | 0.0610870118462435 | -0.0979106210164925 | 0.0823668888540564 | 0 | 0 |
| elbow23 | PSII | 0.00722222222222224 | 0.00717440581881117 | 0.000516130791366882 | 0.0140520193324578 | -0.0016445406078245 | 0.0160575278593774 | 1 | 0 |
| elbow06 | PSV | 0.00757575757575741 | 0.0076789154043771 | -0.00700545242819516 | 0.0219287145132119 | -0.0114309489163429 | 0.0265140463143367 | 0 | 0 |
| elbow28 | PSIII | 0.00757575757575761 | 0.00759718469460701 | -0.00725558864863004 | 0.0220357521799915 | -0.0118112723085827 | 0.0266437006960717 | 0 | 0 |
| elbow04 | PSV | -0.00781250000000008 | -0.00781228396854854 | -0.0222575258392644 | 0.0064814808940771 | -0.0262614486071374 | 0.0110870123728287 | 0 | 0 |
| elbow27 | PSII | -0.00814241571225679 | -0.00832050275373305 | -0.0480429402529088 | 0.0324410436971774 | -0.059542073413319 | 0.0450933482887677 | 0 | 0 |
| elbow29 | PSIb | -0.00818278943278941 | -0.00809318043639618 | -0.0422904354333266 | 0.0265947676858421 | -0.0530402118191731 | 0.0376806575123965 | 0 | 0 |
| elbow18 | PSIa | 0.00884905184338435 | 0.00903134523854146 | -0.031600363376878 | 0.0483439144187124 | -0.045005050674939 | 0.0608900027626782 | 0 | 0 |
| elbow16 | PSV | -0.0089503910068426 | -0.00898964789981955 | -0.0279747631237508 | 0.010391195150418 | -0.0345107952126809 | 0.0164256902279455 | 0 | 0 |
| elbow32 | PSIII | 0.00909090909090909 | 0.00920162333558235 | -0.00819971557748205 | 0.0263120711634111 | -0.0140730374614541 | 0.0322334452967591 | 0 | 0 |
| elbow27 | PSV | 0.00926332382424696 | 0.00959820199502028 | -0.0397719226201089 | 0.0577952222319576 | -0.0568832813207396 | 0.0727676045442617 | 0 | 0 |
| elbow31 | PSIII | -0.00926540037502602 | -0.00913935962521129 | -0.0564298249821211 | 0.0375617441816082 | -0.0713654951526476 | 0.0531457268098275 | 0 | 0 |
| elbow28 | PSV | 0.00940031530439004 | 0.00956964673582372 | -0.0249812856121887 | 0.0429617335325242 | -0.0345401726016496 | 0.0543595866305961 | 0 | 0 |
| elbow02 | PSIII | -0.00984848484848486 | -0.0102079691977347 | -0.0637029535990015 | 0.0460681108756409 | -0.0806668772346348 | 0.0615139412881279 | 0 | 0 |
| elbow20 | PSV | -0.0099730399714594 | -0.00900700389906101 | -0.159303969404843 | 0.134954908984162 | -0.206853502073285 | 0.18142783924479 | 0 | 0 |

| | | | | | | | | | |
|---|---|---|---|---|---|---|---|---|---|
| elbow15 | PSIa | 0.0100509340483743 | 0.0101020859302986 | 0.00217573646574286 | 0.0180325952735333 | -0.000325854210878307 | 0.0204782127319348 | 1 | 0 |
| elbow16 | PSIa | 0.0113207607942007 | 0.0114959014827861 | -0.00362505517379573 | 0.025927695407413 | -0.00812235149611184 | 0.030243507797159 | 0 | 0 |
| elbow06 | PSIII | 0.0113636363636364 | 0.0114429980088675 | -0.0108320484703715 | 0.0331426153084382 | -0.0189315999138023 | 0.0399211324997518 | 0 | 0 |
| elbow14 | PSIb | 0.0116792929292929 | 0.0114916244905993 | -0.0184407497510255 | 0.0413060230918711 | -0.0270967777985296 | 0.0496319077429995 | 0 | 0 |
| elbow08 | PSIII | 0.0117079889807163 | 0.0117773928125343 | -0.00439574582975589 | 0.0277774993261194 | -0.00922522462209594 | 0.0328616750649626 | 0 | 0 |
| elbow16 | PSII | -0.0120059446136001 | -0.0120088459508138 | -0.0331449004112008 | 0.0093912351170274 | -0.0397354868216515 | 0.0162496604414524 | 0 | 0 |
| elbow04 | PSIb | 0.012310606060606 | 0.0119820737882075 | -0.0352826346127417 | 0.0607465167787233 | -0.0474840770541834 | 0.0758194306034998 | 0 | 0 |
| elbow08 | PSII | 0.0123295821825234 | 0.012253257364117 | -0.00968110852843805 | 0.034995043442348 | -0.0161165523166152 | 0.0424505201137123 | 0 | 0 |
| elbow16 | PSIII | -0.0125095492742552 | -0.0127678864366297 | -0.0468928912265474 | 0.021981495087175 | -0.0583887088824994 | 0.0315585321589158 | 0 | 0 |
| elbow08 | PSIa | 0.013035763035763 | 0.0130589134869979 | -0.00450689190283913 | 0.0306444065745184 | -0.00997954164359583 | 0.0359675569717274 | 0 | 0 |
| elbow12 | PSIb | 0.0132575757575757 | 0.0133247709220772 | -0.00437941138382035 | 0.0310479533587282 | -0.0102687293578308 | 0.0369266112382182 | 0 | 0 |
| elbow29 | PSIa | 0.0132656778524677 | 0.0135741898428905 | -0.0312927975336803 | 0.0569633618078306 | -0.043583922352273 | 0.0704045283048104 | 0 | 0 |
| elbow32 | PSIb | -0.0138888888888889 | -0.0137148796229482 | -0.033022658450835 | 0.00551974792617395 | -0.0388878597058413 | 0.0118074265691454 | 0 | 0 |
| elbow07 | PSII | 0.0141754915775961 | 0.014063885632385 | 0.000538846321955205 | 0.0280206962404282 | -0.0034967068504453 | 0.0324314539589351 | 1 | 0 |
| elbow32 | PSII | -0.0150282173966384 | -0.0149431816022926 | -0.0524512662179059 | 0.0217931035955608 | -0.0621048336402045 | 0.0330661336095266 | 0 | 0 |
| elbow04 | PSII | -0.0150421508316246 | -0.0151360546996404 | -0.0510642721504587 | 0.0219385618921792 | -0.0624061886199936 | 0.0323607911539556 | 0 | 0 |
| elbow19 | PSIII | 0.0151515151515152 | 0.0151634200252882 | -0.0139619323514568 | 0.0437097238917713 | -0.0235349311809865 | 0.0532671226776654 | 0 | 0 |
| elbow25 | PSIII | 0.0151515151515152 | 0.015127637055988 | -0.0139446430937931 | 0.0439083716820574 | -0.0221382374054423 | 0.0534018823488622 | 0 | 0 |
| elbow26 | PSII | 0.0152325027444642 | 0.0151187294060425 | 0.00304147805059231 | 0.0275439109875327 | -0.000759946634369676 | 0.031297689693736 | 1 | 0 |
| elbow31 | PSIb | 0.0161227661227662 | 0.0159363578802243 | -0.0348504371857421 | 0.0686469139845903 | -0.0515569695666345 | 0.0858327600173688 | 0 | 0 |
| elbow05 | PSV | -0.0162878787878788 | -0.0161642698255927 | -0.0559077010673077 | 0.0232562511056772 | -0.0676334871476231 | 0.0347908677460851 | 0 | 0 |
| elbow25 | PSIb | -0.0183080808080809 | -0.0176863656866609 | -0.0651838035943434 | 0.0267539330979035 | -0.0781842406074495 | 0.0399173536452364 | 0 | 0 |
| elbow22 | PSV | -0.0183841888194894 | -0.0178706983602927 | -0.0785945588818943 | 0.0419007264514471 | -0.0967075109533518 | 0.0596293218177502 | 0 | 0 |
| elbow01 | PSV | -0.018767217630854 | -0.018785790323049 | -0.0567974405461299 | 0.019886686611676 | -0.0680191097818442 | 0.0326258477912702 | 0 | 0 |
| elbow15 | PSIII | 0.018904958677686 | 0.0188167340076127 | -0.0023512093166005 | 0.0401895003178492 | -0.00787550827247413 | 0.0463398215560841 | 0 | 0 |
| elbow14 | PSIII | 0.0191558441558442 | 0.0192646071527299 | -0.00386562080093405 | 0.0421403722696757 | -0.0119614218454937 | 0.0496237310925823 | 0 | 0 |
| elbow12 | PSII | 0.0194434039958337 | 0.019457364905652 | 0.00296285831285668 | 0.0359013836159515 | -0.00212227750135941 | 0.0411419331497941 | 1 | 0 |
| elbow29 | PSIII | -0.0197314049586776 | -0.0198418508242925 | -0.108760023094804 | 0.0695912803165046 | -0.135177556469368 | 0.0991499228936913 | 0 | 0 |
| elbow02 | PSIb | -0.0198378010878011 | -0.0199239295815698 | -0.0636510364996723 | 0.0243190858237554 | -0.0774231778552472 | 0.0383788917559244 | 0 | 0 |
| elbow17 | PSIa | -0.0202274212512638 | -0.0202903992273477 | -0.081375124624074 | 0.0419693757251012 | -0.101062734068124 | 0.0630050361725613 | 0 | 0 |
| elbow13 | PSV | 0.0203389891645388 | 0.0200673178330646 | -0.0308866840708712 | 0.0728813111264476 | -0.0459191344509512 | 0.0874230496358414 | 0 | 0 |
| elbow24 | PSIb | 0.0205419580419582 | 0.0205523994322303 | -0.0485861046112491 | 0.0902344926827892 | -0.0690114638630405 | 0.113520090402148 | 0 | 0 |
| elbow01 | PSIII | -0.0208333333333333 | -0.0207501080275584 | -0.0594724009894258 | 0.0178001489848489 | -0.0712180753530306 | 0.0302647580928263 | 0 | 0 |
| elbow23 | PSV | -0.0212609970674487 | -0.0215747026707289 | -0.0625786883752324 | 0.0205093699130561 | -0.0745641628463315 | 0.0323111214867129 | 0 | 0 |
| elbow13 | PSIII | -0.0226948630691946 | -0.022517622536326 | -0.0866030488365028 | 0.0407030484353799 | -0.105065299450094 | 0.0605168032755831 | 0 | 0 |
| elbow23 | PSIII | -0.0227272727272727 | -0.022113631303798 | -0.104283745365882 | 0.0567871946658009 | -0.130533538091497 | 0.0841123700897731 | 0 | 0 |
| elbow28 | PSIa | -0.0227652915174628 | -0.0225533407608992 | -0.119732958403972 | 0.0735970314359325 | -0.153071566558143 | 0.101450968368973 | 0 | 0 |
| elbow26 | PSIa | 0.0233584429562743 | 0.0233645619142245 | 0.00321262668516409 | 0.0430822419396865 | -0.00212197839025583 | 0.0492643177274281 | 1 | 0 |
| elbow30 | PSV | -0.023615225441414 | -0.0236850437457532 | -0.0621556993615516 | 0.0160413966206164 | -0.074827109267484 | 0.0287062074040567 | 0 | 0 |
| elbow22 | PSIb | -0.0243055555555556 | -0.0247575896082529 | -0.0855775859770814 | 0.0365003149433325 | -0.103327400861344 | 0.0539526434515306 | 0 | 0 |
| elbow04 | PSIII | -0.025 | -0.0253020678358841 | -0.0710250738153131 | 0.0207631064281267 | -0.0861416727163214 | 0.0347999905116192 | 0 | 0 |
| elbow26 | PSIII | 0.0259740259740257 | 0.0259616564684058 | -0.0035789351410119 | 0.0565670744708047 | -0.0131219741717421 | 0.0651314989703958 | 0 | 0 |
| elbow30 | PSIb | 0.0261606449106449 | 0.0255412398203124 | -0.0456608257213078 | 0.0975436006323735 | -0.0677385293494299 | 0.119599004104006 | 0 | 0 |
| elbow03 | PSII | -0.0272760346614337 | -0.0269971817943186 | -0.0946412610742562 | 0.0389318134875973 | -0.117225258051896 | 0.0582634121900053 | 0 | 0 |
| elbow12 | PSIa | 0.0275887362631695 | 0.0275793492567401 | -0.00259031784936769 | 0.0583493300297583 | -0.0118788892980751 | 0.0668653671295504 | 0 | 0 |
| elbow27 | PSIb | 0.0277777777777777 | 0.0279544913231194 | -0.0165389531059476 | 0.0708304723439572 | -0.0304138309644926 | 0.0851557712899998 | 0 | 0 |
| elbow24 | PSII | 0.0282668089235424 | 0.0282734524872374 | 0.00458122212805128 | 0.0521352809852738 | -0.00398846469964789 | 0.0601108059430522 | 1 | 0 |
| elbow27 | PSIa | -0.0307803232431645 | -0.0308637423604223 | -0.128486660889513 | 0.0676261422928788 | -0.159811888998312 | 0.0975962349242502 | 0 | 0 |
| elbow19 | PSIb | 0.0315656565656566 | 0.031886736586328 | -0.0572452236198576 | 0.118849631639955 | -0.0877729429708554 | 0.144284833702755 | 0 | 0 |
| elbow13 | PSIa | 0.0322744440300675 | 0.0323182453099227 | -0.0236832477858196 | 0.0882502270660854 | -0.040894362166181 | 0.10496561361099 | 0 | 0 |
| elbow02 | PSII | -0.0341864305949235 | -0.0341710257297277 | -0.0786662538436544 | 0.0101291627402343 | -0.0930936727131151 | 0.0235775730662045 | 0 | 0 |
| elbow18 | PSV | 0.0352462488003754 | 0.0351738681343908 | 0.000181267821958148 | 0.0701845511442362 | -0.01022791598117 | 0.0816217680730943 | 1 | 0 |
| elbow33 | PSIa | 0.0359986615663316 | 0.0358696793711584 | 0.00482731314708232 | 0.0671763399440162 | -0.00577713585967439 | 0.0768163112976923 | 1 | 0 |
| elbow32 | PSV | -0.0391277673350042 | -0.040142788659445 | -0.107096703583476 | 0.0311622244212954 | -0.129862607495888 | 0.0511491629693295 | 0 | 0 |
| elbow33 | PSII | 0.0396445659603555 | 0.0398522048977806 | -0.0332662982781187 | 0.112082062398873 | -0.0575504939529457 | 0.13417394677016 | 0 | 0 |
| elbow09 | PSIII | -0.0398268398268398 | -0.0397674070663457 | -0.0932709955061425 | 0.0144530925346463 | -0.109408113123053 | 0.0310825630934072 | 0 | 0 |
| elbow20 | PSII | 0.0438799597652699 | 0.0429996791389032 | -0.0384685367756634 | 0.128494098633033 | -0.0635422198404513 | 0.155420849077802 | 0 | 0 |
| elbow24 | PSV | 0.044290474992778 | 0.0439282693105675 | -0.0279672782636496 | 0.115487367120846 | -0.0490631599447994 | 0.140730294161557 | 0 | 0 |
| elbow13 | PSII | 0.0449175927649822 | 0.0450361991168396 | 0.00219346230204555 | 0.0855437324448897 | -0.0100114770758752 | 0.0999774422699198 | 1 | 0 |
| elbow10 | PSIb | 0.0482566045066045 | 0.047753871976538 | -0.0300010503507835 | 0.126200996880541 | -0.0544698466990553 | 0.150135727859157 | 0 | 0 |
| elbow08 | PSIb | 0.048951048951049 | 0.0479640393106262 | -0.0336861052903406 | 0.135523406883805 | -0.0630203799710996 | 0.163164212796809 | 0 | 0 |
| elbow02 | PSV | -0.0504545454545455 | -0.0503689363649102 | -0.133923703171053 | 0.0324799521677665 | -0.161533003444324 | 0.0589849567499375 | 0 | 0 |
| elbow33 | PSIb | 0.0505050505050505 | 0.0499305652003921 | -0.0348666450954271 | 0.138079474703876 | -0.0656403702065573 | 0.164606877024563 | 0 | 0 |
| elbow13 | PSIb | 0.0505293317793318 | 0.0505090958224124 | -0.00900449582542516 | 0.110183666629398 | -0.0253899902452369 | 0.128174002798385 | 0 | 0 |
| elbow21 | PSIb | 0.0514374514374515 | 0.0507227197551403 | -0.0859270979687797 | 0.194433235520456 | -0.134681338706186 | 0.238197275418701 | 0 | 0 |
| elbow25 | PSV | -0.0523735733963007 | -0.0520805162556627 | -0.112766502344584 | 0.00918977667694684 | -0.133543905928864 | 0.0260883906480636 | 0 | 0 |
| elbow21 | PSIII | 0.0544103178461468 | 0.0543980936953206 | -0.000198112581770682 | 0.110336409082272 | -0.018285351049264 | 0.128419840102827 | 0 | 0 |
| elbow26 | PSIb | -0.0565656565656565 | -0.0566467258436847 | -0.233356978155262 | 0.115227502625693 | -0.284095661291464 | 0.171313025856074 | 0 | 0 |
| elbow05 | PSIa | -0.0591040716040717 | -0.0595623071551925 | -0.174016027344068 | 0.0551920696020131 | -0.212795550874357 | 0.0921487708545622 | 0 | 0 |
| elbow27 | PSIII | -0.0600229761326019 | -0.059786943739875 | -0.16398124450224 | 0.0416143453925644 | -0.195048243324607 | 0.0737215001908556 | 0 | 0 |
| elbow07 | PSIb | -0.0624999999999999 | -0.0619475341587087 | -0.177766354992006 | 0.0499006060633896 | -0.218341710888638 | 0.0896303980231575 | 0 | 0 |
| elbow21 | PSIa | 0.0666156498575894 | 0.0660729959633815 | -0.00798397017027392 | 0.143641400323724 | -0.0347378385583422 | 0.166164609326389 | 0 | 0 |
| elbow10 | PSIII | 0.0671733569460844 | 0.0668270668874155 | -0.0357125137615257 | 0.172389418519694 | -0.0758725398512603 | 0.202674489820909 | 0 | 0 |
| elbow04 | PSIa | -0.0681786735541002 | -0.0679898230317469 | -0.187681793584891 | 0.0503988286080355 | -0.221472161909834 | 0.0834354418198451 | 0 | 0 |
| elbow03 | PSIa | -0.0693575595452732 | -0.0693925230563471 | -0.223357179743501 | 0.0830425238172113 | -0.270753944346763 | 0.130901820418748 | 0 | 0 |
| elbow21 | PSV | 0.0697862039753321 | 0.0697692200519932 | 0.0145767218494161 | 0.124956719724361 | -0.00103836205180636 | 0.141846328983436 | 1 | 0 |
| elbow18 | PSIII | 0.081147128272797 | 0.0806424493357246 | -0.00982133801625554 | 0.175464276324435 | -0.0402745971506599 | 0.203996533299361 | 0 | 0 |
| elbow29 | PSV | -0.0820922478562204 | -0.0823268242557697 | -0.1731318167531 | 0.00954947749568932 | -0.202781919562713 | 0.0380569631408318 | 0 | 0 |
| elbow10 | PSIa | -0.0901265118914986 | -0.0907986830296714 | -0.200277357065606 | 0.0193614064810138 | -0.23400762592694 | 0.0556247160396554 | 0 | 0 |
| elbow20 | PSIb | 0.0918803418803421 | 0.0915975360296477 | -0.0321694807266479 | 0.215824316618756 | -0.0677184013607661 | 0.252804874960677 | 0 | 0 |
| elbow02 | PSIa | -0.0981180466589231 | -0.0982931046889943 | -0.27073150032894 | 0.0723806980737637 | -0.325868399231424 | 0.124247682125091 | 0 | 0 |
| elbow09 | PSIa | -0.107702285389999 | -0.107131436862043 | -0.260491647639119 | 0.0414555935943426 | -0.306882841202082 | 0.0856793959346072 | 0 | 0 |
| elbow24 | PSIII | 0.121096661805218 | 0.121062630324984 | 0.00483679976342845 | 0.240106751098699 | -0.0334436520764023 | 0.277252922575671 | 1 | 0 |

**Supplementary table 6. Statistical results from linear regression model bootstrapping for GBS phenotype and population analysis.** The observed (obs_estimate) and estimated (unbiased_estimate) difference in proportions of clusters among total B cells for a given serotype-specificity and donor population (South African- over Dutch donors) were generated with a two-sided linear regression model with wild bootstrap simulation (9999 resamples). Confidence intervals (ci) and Bonferroni's method adjusted ci (ci.adj) are shown, as well as statistical significance (signif_ci/ signif_ci.adj = 1).
